# *MONOPTEROS* isoform *MP11ir* role during somatic embryogenesis in *Arabidopsis thaliana*

**DOI:** 10.1101/2024.07.17.603838

**Authors:** Barbara Wójcikowska, Samia Belaidi, Victoria Mironova, Sylvie Citerne, Hélène S. Robert

**Affiliations:** Hormonal Crosstalk in Plant Development, Mendel Centre for Plant Genomics and Proteomics, CEITEC MU - Central European Institute of Technology, Masaryk University, Brno, Czech Republic; Institute of Biology, Biotechnology and Environmental Protection, Faculty of Natural Sciences, University of Silesia in Katowice, Katowice, Poland; Laboratory of Functional Genomics and Proteomics, National Centre for Biomolecular Research, Faculty of Science, Masaryk University, Brno, Czechia; Department of Plant and Animal Biology, Radboud Institute for Biological and Environmental Sciences (RIBES), Radboud University, The Netherlands; Université Paris-Saclay, INRAE, AgroParisTech, Institute Jean-Pierre Bourgin for Plant Sciences (IJPB), 78000, Versailles, France

**Keywords:** Alternative splicing, *Arabidopsis thaliana*, Auxin, AUXIN RESPONSE FACTOR, Auxin signalling, *In vitro* culture, Isoform, MONOPTEROS, MP11ir, Somatic embryogenesis, Transcription factor

## Abstract

Auxin is crucial for plant morphogenesis, including *in vivo* and *in vitro* embryo development. Exogenous auxin application is necessary for inducing embryogenic responses in *in vitro* cultured explants of Arabidopsis and other plants. Thus, components of auxin transport, signaling, and metabolism are key to somatic embryogenesis. AUXIN RESPONSE FACTOR (ARF) transcription factors bind to auxin response elements to control auxin-responsive gene expression and are vital in somatic embryo regeneration. ARFs are often repressed by AUXIN/INDOLE-3-ACETIC ACIDs (Aux/IAAs). MONOPTEROS (MP)/ARF5 is especially important in the embryogenic transition, being highly expressed during somatic embryogenesis; its mutant cannot develop somatic embryos. The *MP11ir* transcript, an alternatively spliced variant of *MP*, produces a truncated protein missing the Phox and Bem1p (PB1) domain, crucial for ARF-Aux/IAA dimerization. This renders the MP11ir isoform insensitive to Aux/IAA repression, suggesting an auxin-independent regulation. High levels of *MP11ir* transcript are observed during auxin- and trichostatin A-dependent induction of somatic embryogenesis. Both MP and MP11ir are essential for embryo regeneration in the *mpS319* mutant. However, overexpression of a truncated MP protein (ΔARF5) lacking the PB1 domain inhibits somatic embryogenesis, resulting in callus formation instead of somatic embryos. Overexpression of Δ*ARF5*, lack of MP protein (*mp* mutant), and interference with MP action by the auxin-resistant BODENLOS (BDL) protein affect the expression of the genes *TRYPTOPHAN AMINOTRANSFERASE OF ARABIDOPSIS 1* (*TAA1*), *TAA1-RELATED 1* (*TAR1*), *YUCCA3* (*YUC3*), *YUC5* and *YUC8*, which are involved in auxin biosynthesis. The partial complementation of *mpS319* by the expression of *MP11ir* may be due to the restoration of the expression of *TAA1*, *TAR1*, *YUC3, YUC5,* and *YUC8* to the levels observed in the wild-type genotype Col-0. The presence of MP protein alone or in combination with MP11ir in *mpS319* restored the expression of all analyzed genes compared to Col-0. Our results suggest that the analyzed auxin-related genes could be potential targets of MP11ir and/or MP. Consequently, Δ*ARF5* overexpression alters auxin homeostasis and endogenous auxin levels, hindering embryogenic transition.

## Introduction

Auxin plays a vital role in various aspects of plant development, including plant growth and morphogenesis in response to environmental signals through its transport, metabolism, and signaling (Caumon and Vernoux, 2023; Cohen and Strader, 2024; Liu et al., 2024). Recognized as a key trigger of developmental changes, auxin is a critical inducer of somatic embryogenesis (SE) in many plant species (Elhiti and Stasolla, 2022). SE is a unique plant-specific developmental process that demonstrates the pluripotency of somatic cells and is widely exploited for *in vitro* plant regeneration. This technique is used in biotechnology for mass micropropagation of plants (Mehbub et al., 2022), plant biodiversity conservation (Ballesteros et al., 2024), and the production of transgenic plants (Lee and Wang, 2023; McFarland and Kaeppler, 2024; Yang et al., 2024). With the rising global population, decreased crop yields due to climate change, and increasing environmental pollution, the importance of *in vitro* plant propagation techniques is growing (Hasnain et al., 2022; Krasteva et al., 2021). Understanding the molecular mechanisms of SE can enhance its efficiency and its optimization for recalcitrant species (Chen et al., 2022). Exogenous auxin treatment of explants modifies chromatin accessibility and the transcriptome of somatic cells (Wang et al., 2020; Wu et al., 2022). This modulation affects many genes, including those encoding transcription factors (TFs), proteins associated with hormone transport, metabolism, signaling, and stress response. The induction of SE is also associated with the expression of the genes encoding the core components of the nuclear auxin signaling pathway, such as AUXIN RESPONSE FACTORs (ARFs) and AUXIN/INDOLE-3-ACETIC ACIDs (Aux/IAAs) (Quintana-Escobar et al., 2019). Thus, the crosstalk between auxin and these transcriptional regulators is fundamental to the SE regulatory network.

MONOPTEROS (MP), known as AUXIN RESPONSE FACTOR 5 (ARF5), is pivotal in nuclear auxin signaling and has been identified in many plant species (Chandler, 2016). The function of MP is well-characterized, thanks to a wide range of tools like allelic mutants, overexpressing and reporter lines in *Arabidopsis thaliana,* allowing for comprehensive functional, genetic, and microscopic analyses (Hardtke et al., 2004; Odat et al., 2014; Rademacher et al., 2011; Schlereth et al., 2010; Wójcikowska et al., 2023; Wu et al., 2015). Knowledge of the MP action is being transferred to commercially important species (Liu et al., 2018a; Xu et al., 2019; Yue et al., 2020). MP is crucial for many aspects of *in vivo* plant development, including flower, ovule, and pollen development (Cucinotta et al., 2021; Liu et al., 2018b; Wu et al., 2015), zygotic embryogenesis (Möller et al., 2017), root and shoot meristem establishment (Cole et al., 2009; Krogan et al., 2016; Schlereth et al., 2010), organ polarity maintenance (Bhatia et al., 2016), and vascularization patterns (Mattsson et al., 2003; Przemeck et al., 1996). MP is also critical for *in vitro* plant morphogenesis. *MP* is essential for an effective SE process because its expression is stimulated during SE; it is active in regions (cotyledons and SAM) where somatic embryos emerge; and, more importantly, explants of *mp* mutant cannot produce somatic embryos (Wójcikowska and Gaj, 2017).

The MP protein has four distinct domains: a B3-type DNA binding domain (DBD), dimerization domains (DD1, DD2), a middle region (MR) region, and a Phox and Bem1 (PB1) domain (Guilfoyle, 2015). MP binds to the AuxRE *cis*-elements in the promoter of target genes. As a transcription factor, MP primarily acts as a positive regulator of gene expression (Boer et al., 2014; Freire-Rios et al., 2020). In Arabidopsis, MP binds as a dimer to AuxRE motifs organized as inverted repeats (IR), direct repeats (DR), and everted repeats (ER). IR consists of two AuxREs on opposite DNA strands facing each other, DR has two consecutive AuxREs on the same DNA strand, and ER features two AuxREs on opposite directions and DNA strands. The DD1 and DD2 domains flank the DBD, acting as a caliper for binding to IR motifs. For docking on ER and DR motifs, the MP DBD dimer relies on the MR or PB1 domains (Cancé et al., 2022). Additionally, AuxRE elements in the MP promoter indicate that its expression is regulated by cellular auxin level (Lau et al., 2011).

Three scenarios outline MP action. First, at low auxin levels, MP interacts with BODENLOS/INDOLE-3-ACETIC ACID 12 (BDL/IAA12) via PB1 domains, recruiting TOPLESS/TOPLESS RELATED (TPL/TPR), transcriptional co-repressor proteins, and HDA19 histone deacetylase enzyme. HDA19 removes the acetyl group from the histone H3 and H4 tails, condensing the chromatin and blocking the expression of auxin-responsive genes (Chandler, 2016). In the presence of auxin, BDL is recruited by the SCF^TIR/AFBs^ complex, ubiquitinated, and degraded by the proteasome, releasing MP to form a complex with SWI/SNF chromatin remodeling ATPases BRAHMA (BRM) and SPLAYED (SYD). It unlocks the chromatin to provide histone acetyltransferases (HATs) and transcription factors access to the *cis-*elements of MP-regulated promoters, thus regulating gene transcription (Wu et al., 2015). Second, above-threshold auxin levels may potentially cause MP protein oligomerization with itself or other ARFs, inhibiting the expression of auxin-responsive genes, as observed *in vitro* (Korasick et al., 2014; Nanao et al., 2014) and *in vivo* for ARF19 (Powers et al., 2019). This scenario is plausible as plants with *MP* constitutive overexpression (Hardtke et al., 2004) phenocopy *mp* mutants (Przemeck et al., 1996). Third, the MP11ir isoform, produced by alternative splicing (AS) of the *MP* transcript during ovule and root development, lacks the PB1 domain due to the retention of intron 11, resulting in the translation of a truncated protein. This isoform is insensitive to Aux/IAA repression and could function independently of auxin. Ectopic expression of *MP11ir* partially complements *mp* mutant phenotypes during reproductive development, indicating that some MP functions do not require interaction with Aux/IAA repressors. MP11ir activates target genes in ovule and root regions with low and high auxin concentrations (Cucinotta et al., 2021; Cavalleri et al., 2024).

To understand the role of MP11ir in the embryogenic transition, we analyzed its transcript level in Arabidopsis explants cultured *in vitro* and treated with SE inducers: auxins 2,4-dichlorophenoxyacetic acid (2,4-D), 1-naphthaleneacetic acid (NAA), and indole-3-acetic acid (IAA), and trichostatin A (TSA). High *MP11ir* transcript levels were associated with auxin- and TSA-dependent SE inductions. MP11ir can partially rescue the *mp* phenotype, improving its capacity for embryogenic transition. However, both MP and MP11ir are required to fully complement the *mp* phenotype. Overexpression of the truncated MP protein (ΔARF5) inhibits SE and alters the expression of auxin biosynthetic genes of the indole-3-pyruvic acid (IPyA) pathway: *TRYPTOPHAN AMINOTRANSFERASE OF ARABIDOPSIS 1* (*TAA1*), *TAA1-RELATED 1* (*TAR1*), *YUCCA 3* (*YUC3*), *YUC5,* and *YUC8*. This modulation changes endogenous auxin levels, negatively impacting embryogenic transition. We hypothesized that MP11ir may directly control *TAA1, YUC3, YUC5,* and *YUC8*. These findings suggest that MP11ir is involved in SE induction alongside the canonical MP, contributing to the regulation of auxin biosynthesis and homeostasis during this process.

## Results

### The presence of *MP11ir* transcript is specific for auxin-dependent embryogenic transition

*MP11ir* was first detected in ovules, where it likely regulates target genes in cells with low auxin concentrations (Cucinotta et al., 2021). Given the critical role of auxin signaling proteins in the embryogenic transitions, we investigated whether *MP11ir* would be active during SE induced by exogenously applied synthetic auxin 2,4-D. We hypothesized that if *MP11ir* regulates its target genes in cells with low auxin concentrations, its presence would be minimal in cells undergoing embryogenic reprogramming, as those cells accumulate exogenous 2,4-D and endogenous IAA (Schulz and Segobye, 2016; Wójcikowska and Gaj, 2015). To test this, we analyzed *MP11ir* transcript levels in various samples: 10-day-old seedlings, flowers, leaves, immature zygotic embryos (IZEs), and IZE explants during auxin-dependent SE induction. *MP11ir* was not detected in leaves but was present in other tissues, notably flowers, as previously observed (Fig. S1) (Cucinotta et al., 2021). Quantitative analysis revealed that *MP11ir* was most abundant during the auxin-dependent embryogenic transition, with 11-18 times higher expression than in seedlings (Fig. 1A). It suggests that *MP11ir* may regulate target gene expression in both low (Cucinotta et al., 2021) and high auxin concentrations (our results; Cavalleri et al., 2024), indicating a role in auxin-independent transcription control during SE.

**Figure 1.**
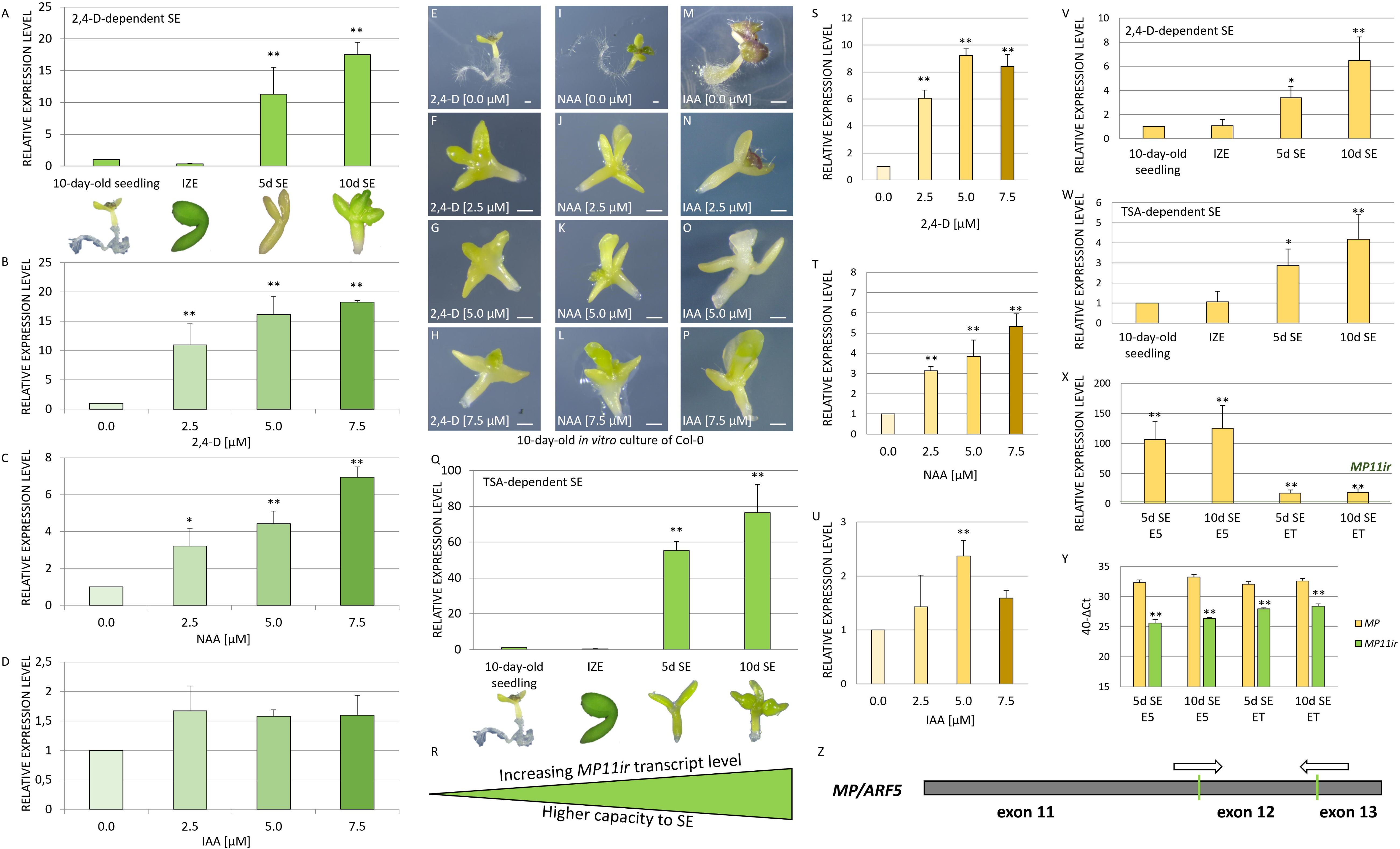
*MP11ir* and *MP* are expressed during somatic embryogenesis induction. (**A**) The level of *MP11ir* transcript in IZE explants and during auxin-dependent (5.0 µM 2,4-D) SE induction (5^th^ and 10^th^ day of *in vitro* culture). Values significantly different from the control culture (10-day-old seedlings) are marked with asterisks. The relative expression level was normalized to internal control (At4g27090) and calibrated to the 10-day-old seedling. A two-way ANOVA analysis (*p* < 0.05) followed by Tukey’s HSD was used to determine any values that were significantly different from 10-day-old seedlings (\**p* < 0.05; \*\**p* < 0.01) (n = 3; means ± SD are presented). (**B-P**) The level of *MP11ir* transcript in the IZE explants cultured for ten days on medium without exogenous auxin, control condition (**E, I, M**), and treated with 2,4-D (**B**), NAA (**C**) and IAA (**D**) concentrations of 2.5 (**F, J, N**), 5.0 (**G, K, O**), and 7.5 µM (**H, L, P**). Values significantly different from the control culture (explants untreated with 2,4-D, NAA, or IAA) are marked with asterisks. The relative expression level was calibrated to the 10-day-old IZE control culture untreated with exogenous auxin. A two-way ANOVA analysis (*p* < 0.05) followed by Tukey’s HSD was used to determine any values that were significantly different from untreated IZE explants (\**p* < 0.05; \*\**p* < 0.01) (n = 3; means ± SD are presented). Size bars indicate 1 mm (**E-P**). (**Q**) The *MP11ir* transcript level in IZE explants and during TSA-dependent (1.0 µM TSA) SE induction (5^th^ and 10^th^ day of *in vitro* culture). Values significantly different from the control culture (10-day-old seedlings) are marked with asterisks. A two-way ANOVA analysis (*p* < 0.05) followed by Tukey’s HSD was used to determine any values that were significantly different from 10-day-old seedlings (\**p* < 0.05; \*\**p* < 0.01) (n = 3; means ± SD are presented). (**R**) Together with data presented in Supplementary Figure S2, we hypothesize a positive correlation between *MP11ir* levels and SE capacity. (**S-U**) The level of *MP* transcript in the IZE explants cultured on medium without exogenous auxin and treated with 2,4-D (**S**), NAA (**T**), and IAA (**U**) concentrations of 2.5, 5.0, and 7.5 µM. Values significantly different from the control culture (explants untreated with 2,4-D, NAA, or IAA) are marked with asterisks. The relative expression level was calibrated to the 10-day-old IZE control culture untreated with exogenous auxin. A two-way ANOVA analysis (*p* < 0.05) followed by Tukey’s HSD was used to determine any values that were significantly different from untreated IZE explants (\**p* < 0.05; \*\**p* < 0.01) (n ≥ 3; means ± SD are presented). (**V, W**) The expression profile of *MP* in IZE explants and during auxin-dependent (5.0 µM 2,4-D) (**V**) and TSA-dependent (1.0 µM TSA) (**W**) SE induction (5^th^ and 10^th^ days of *in vitro* culture). Values significantly different from the expression 10-day-old seedlings are marked with asterisks. A two-way ANOVA analysis (*p* < 0.05) followed by Tukey’s HSD was used to determine any values that were significantly different from 10-day-old seedlings (\**p* < 0.05; \*\**p* < 0.01) (n ≥ 3; means ± SD are presented). (**X**) The expression profile of *MP* during auxin-dependent (E5) and TSA-dependent (ET) SE induction (5^th^ and 10^th^ days of *in vitro* culture). The relative expression level was calibrated to the *MP11ir* expression on the same day of *in vitro* culture. Values significantly different from the *MP11ir* expression are marked with asterisks. A two-way ANOVA analysis (*p* < 0.05) followed by Tukey’s HSD was used to determine any values that were significantly different from 10-day-old seedlings (\**p* < 0.05; \*\**p* < 0.01) (n ≥ 3; means ± SD are presented). (**Y**) The transcript level of *MP* and *MP11ir* during the auxin (E5) and TSA-dependent (ET) SE process (5^th^ and 10^th^ days of *in vitro* culture). The expression level was calculated using the 40-ΔCt method. The At4g27090 gene was used as internal control and for normalization. Values significantly different from the expression of *MP* are marked with asterisks. A two-way ANOVA analysis (*p* < 0.05) followed by Tukey’s HSD was used to determine any values that were significantly different from *MP* expression (\**p* < 0.05; \*\**p* < 0.01) (n ≥ 3; means ± SD are presented). (**Z**) The location of primers used to detect *MP* transcript is indicated.

### The presence of the *MP11ir* transcript depends on the type and concentration of exogenously applied auxin

To investigate the correlation between *MP11ir* transcript levels and auxin presence, we treated IZEs with various auxins, i.e., 2,4-D, NAA, and IAA, at increasing concentrations (0, 2.5, 5.0, and 7.5 µM) for ten days. Different auxins at the same concentrations had varied effects on *MP11ir* levels. We observed a positive correlation between *MP11ir* levels and increasing concentrations of 2,4-D (Spearman correlation coefficient 0.87; p < 0.01), NAA (Spearman correlation coefficient 0.92; p < 0.01), but not IAA (Fig. 1 B-D). The application of 2,4-D had the most substantial promoting effect on *MP11ir* levels. A 2.5 µM concentration of 2,4-D increased *MP11ir* transcript levels 10-fold compared to untreated explants, while a 7.5 µM concentration of 2,4-D (three times higher) resulted in an 18-fold increase (Fig. 1B, E-H). The highest concentration of NAA (7.5 µM) only increased *MP11ir* levels sevenfold (Fig. 1C, I-L). IAA did not significantly affect MP11ir transcript levels (Fig. 1D, M-P). The strength of SE induction also varied with the type of applied auxins (Fig. S2). In Arabidopsis IZE culture, 2,4-D is the most effective for SE induction (Fig. S2A, D-G), followed by NAA (Fig. S2B, H-K), with IAA being the least effective (Fig. S2C, L-O). We found a positive correlation between the increase of *MP11ir* transcript levels and the strength of SE induction for 2,4-D (Spearman correlation coefficient 0.80; p < 0,01) and NAA (Spearman correlation coefficient 0.75; p < 0,01) but not for IAA. This data suggests that the type and concentration of auxin influence *MP11ir* levels and subsequently affect the efficiency of the SE process.

### The presence of *MP11ir* transcript is specific for the embryogenic transition independently of the presence of auxin

Synthetic auxin 2,4-D is the most common and effective inducer of SE in Arabidopsis and other plants (Wójcik et al., 2020). To determine whether *MP11ir* is specific for embryogenic transition rather than merely responding to the application of exogenous auxins, which may also act as stressors (Karami et al., 2023), we assessed the impact of trichostatin A (TSA), an inhibitor of histone deacetylases (HDAC) known to induce SE (Tanaka et al., 2008), on the levels of *MP11ir* transcript. IZE explants treated with 1 µM TSA rapidly and massively regenerate somatic embryos (Wójcikowska et al., 2018). We measured *MP11ir* levels on the 5^th^ and 10^th^ days of TSA-dependent SE induction and found significantly elevated levels of *MP11ir*, 55 and 84 times higher than in seedlings, respectively (Fig. 1Q). These findings suggest that MP11ir is specifically associated with the developmental switch from somatic to embryonic cell fate, highlighting its role in the embryonic transition (Fig. 1R).

### The *MP* transcript is detected during the SE process

The transcript encoding the canonical MP protein was also detected during the SE process, which is consistent with current knowledge (Wójcikowska and Gaj, 2017). Its expression pattern was very similar to that of *MP11ir*. The *MP* transcript significantly accumulated in 2,4-D and NAA-treated IZE explants (all concentrations tested) and in explants treated with 5.0 µM IAA (Fig. 1S-U). We observed a significant up-regulation of *MP* expression during auxin-dependent (Fig. 1V) and TSA-dependent (Fig. 1W) induction of the SE process. During auxin- and TSA-induced SE, *MP* is predominantly synthesized over *MP11ir* (Fig. 1X, Y). During the auxin-induced embryogenic transition, the transcript level of *MP* was more than 100-fold higher than that of *MP11ir*, whereas it was nearly 18-fold higher during TSA-induced SE (Fig. 1X). These results indicate that both *MP* (Fig. 1Z) and *MP11ir* (Fig. S1B) transcripts are synthesized during somatic embryogenesis, denoting their involvement in the embryogenic transition.

### Both MP and MP11ir are necessary to fully rescue the SE-related phenotypes of the *mpS319* mutant

Mutations in the *MP* gene dramatically reduce the capacity of IZE explants for SE. The *mpS319* mutant showed impaired SE efficiency (20%) and productivity (2 somatic embryos per explant) compared to Col-0 (85% efficiency and 4 embryos per explant) (Fig. 2A-D, H-J). Mutations in *MP* affect zygotic embryo development, potentially impacting the capacity of the IZE to undergo SE. To address this, we inhibited the MP-dependent signaling pathway by expressing an auxin-insensitive BDL/IAA12 (bdl) protein in the IZE. Overexpression of DEX-induced *bdl* in IZE completely abolished the embryogenic transition. Explants of the *p35S::bdl-GR* line treated with DEX did not respond to the 2,4-D SE inducer and developed into seedlings. In contrast, Col-0 and the *p35S::BDL-GR* transgenic line expressing a native BDL protein that is ubiquitinated and degraded by the proteasome in the presence of auxin, regenerated somatic embryos massively and independently of the presence of DEX (Fig. 2N-X). These results confirm the findings obtained with the *mpS319* line.

**Figure 2.**
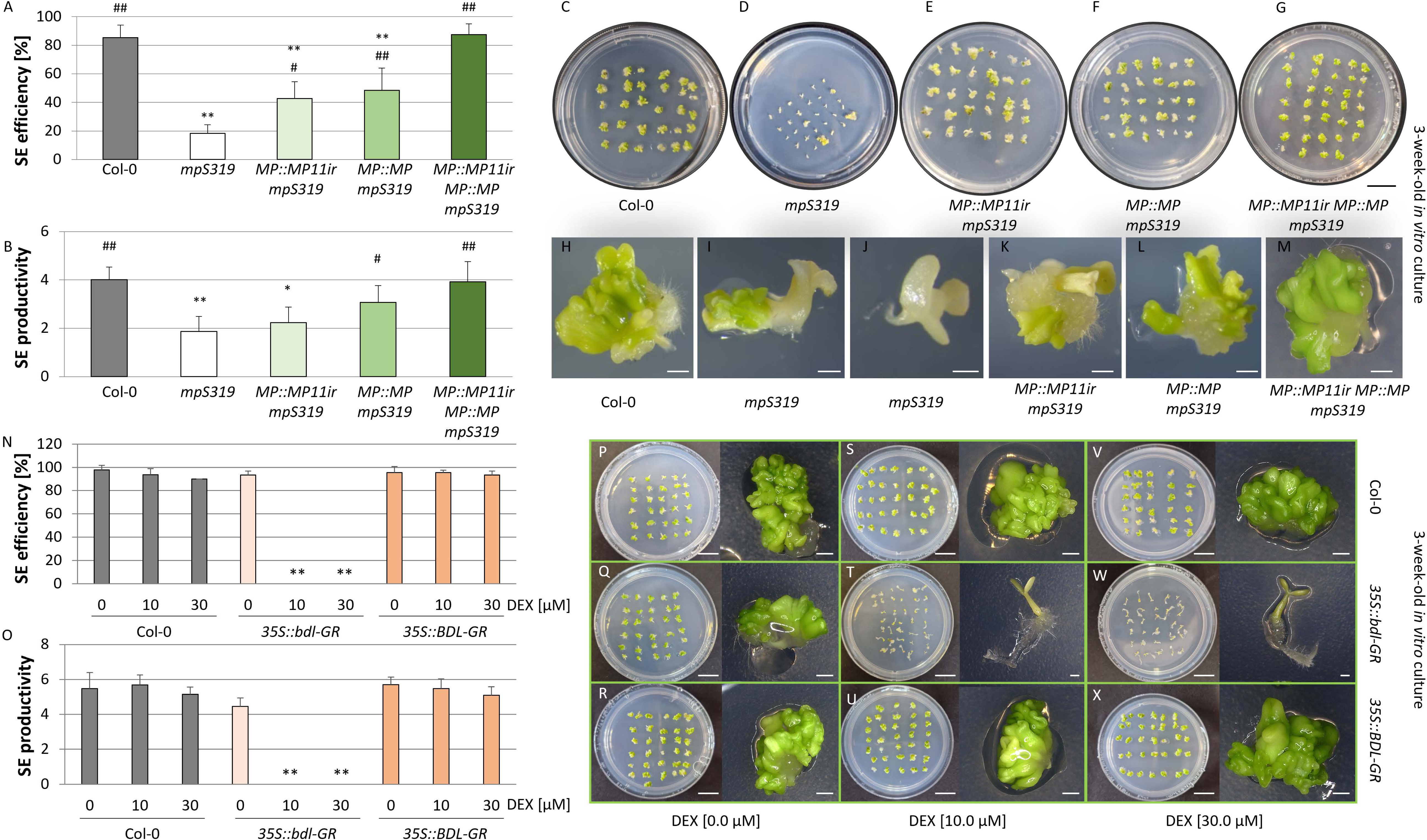
MP and MP11ir are both required to rescue the reduced SE capacity of *mpS319*. (**A-M**) The embryogenic capacity (**A, B**) of the Col-0 (**C, H**), *mpS319* mutant (**D, I, J**), *pMP::MP11ir* (**E, K**), *pMP::MP* (**F, L**) and *pMP::MP11ir pMP::MP* (**G, M**) transgenes in *mpS319* background on the E5 medium evaluated in a 21-day-old culture. Values significantly different from the control culture (Col-0) are marked with asterisks, and from *mpS319* mutant are marked with hashtags. A two-way ANOVA analysis (*p* < 0.05) followed by Tukey’s HSD was used to determine any values that were significantly different (*^, #^*p* < 0.05; **^, # #^*p* < 0.01) (n ≥ 3; means ± SD are given). (**N-X**) The embryogenic capacity (**N, O**) of the Col-0 (**P, S, V**), *p35S::bdl-GR* (**Q, T, W**), *p35S::BDL-GR* (**R, U, X**) transgenic lines on the E5 medium (**P, Q, R**) and E5 medium supplemented with 10 µM (**S, T, U**) and 30 µM (**V, W, X**) of DEX. SE capacity was evaluated in 21-day-old cultures. Values significantly different from the control culture (Col-0) untreated with DEX are marked with asterisks. A two-way ANOVA analysis (*p* < 0.05) followed by Tukey’s HSD was used to determine any values that were significantly different (\**p* < 0.05; \*\**p* < 0.01) (n = 3; means ± SD are presented). Size bars indicate 1 mm for explant images and 1 cm for plate images.

It was previously shown that MP11ir could partially rescue the *mpS319* mutant phenotypes during ovule development (Cucinotta et al., 2021). To investigate the function of MP11ir and its involvement in the SE process, we performed a complementation assay by expressing *MP11ir* or/and *MP* under the control of the *MP* promoter (*pMP::MP11ir-GFP*, hereafter *pMP:MP11ir* and *pMP::MP* transgenes) in the *mpS319* mutant. The presence of MP11ir improved the SE capability of the *mpS319* mutant by 2.3-fold (Fig. 2A-B, E, K). Interestingly, canonical MP did not fully rescue the *mpS319* SE defects, as only 48% of explants responded to SE induction in *MP::MP mpS319* explants (Fig. 2A-B, F, L). Both *MP* and *MP11ir* transcripts must be present to fully rescue the *mpS319* phenotype (Fig. 2A-B, G, M). The complementation assay showed that MP11ir and MP are required together to fully rescue the *mpS319* SE defects and that both the canonical and truncated MP proteins are necessary for effective somatic embryo formation.

### MP and MP11ir localization during the SE process

To detect the MP and MP11ir protein during the auxin-dependent SE process, we used the transgenic lines *pMP::MP-GFP* and *pMP::MP11ir-GFP*. The MP signal was detected at different stages of SE induction (0, 1, 5, 10 days of culture) as a strong nuclear signal in the shoot apical meristem, cotyledons, and root tip (Fig. 3A, C, E, G). Notably, the MP nuclear signal progressively intensified and became widespread as the SE ages. In contrast, only faint and diffuse fluorescence signal was detected for MP11ir at all stages of culture (Fig. 3 B, D, F, H). The signal often lacks a clear nuclear localization pattern as observed for MP-GFP. This suggests that MP11ir protein is either (i) expressed below detection levels, (ii) is not expressed in these tissues under our experimental conditions, or (*iii*) subjected to post-transcriptional or - translational modifications resulting in the absence of a clear nuclear GFP signal. Using available lines, only MP protein was present during the SE process, suggesting that MP and MP11ir might have distinct roles during the SE process.

**Figure 3.**
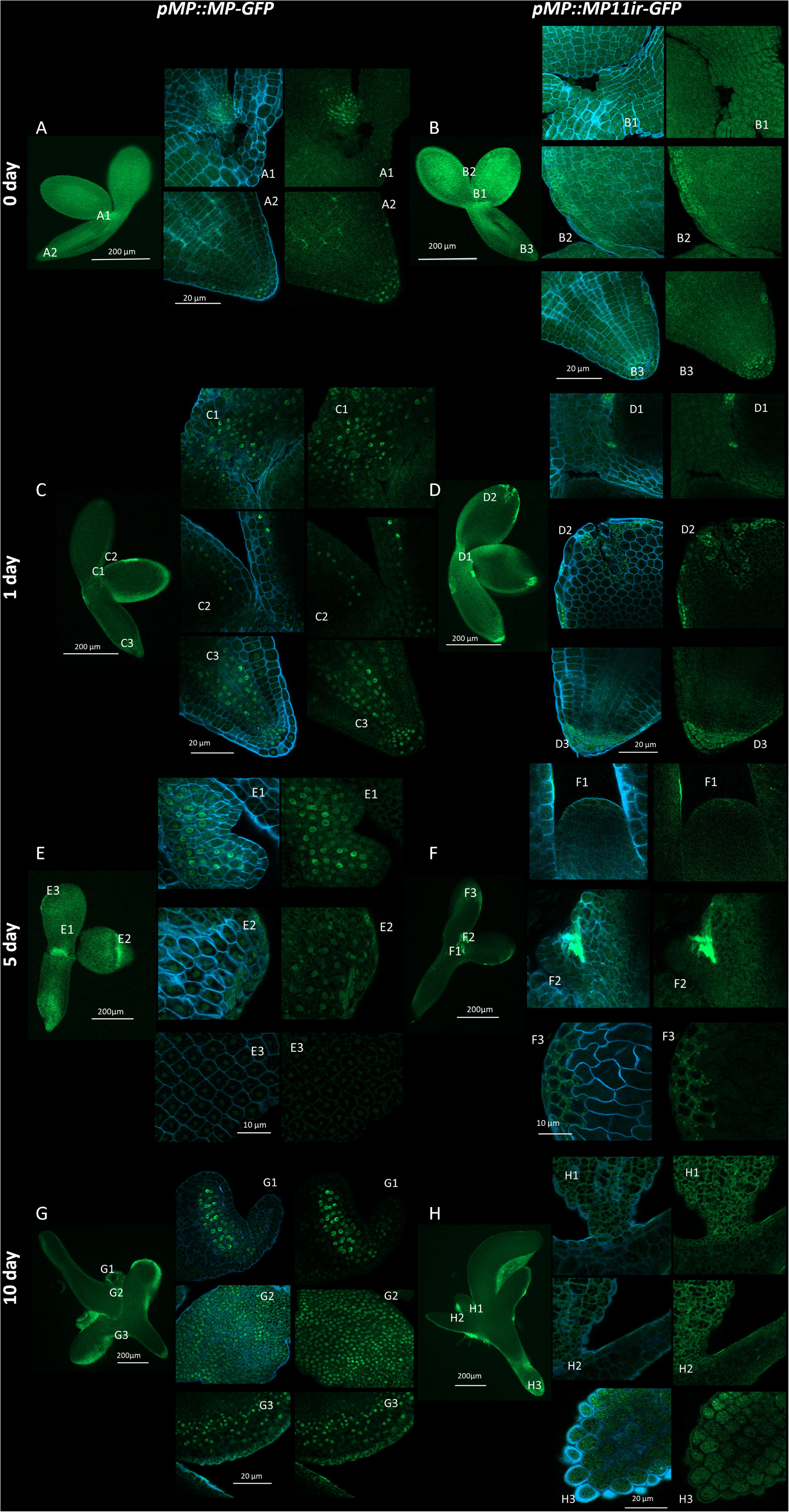
Expression pattern of MP and MP11ir during SE induction. Spatio-temporal localization of MP (**A**) and MP11ir (**B**) in IZE explants and explants cultured on the SE induction medium with 5 μM 2,4-D by 1 (**C**, **D**), 5 (**E**, **F**) and 10 (**G**, **H**) days of SE culture. d, day of SE culture. IZE explants were cultured from *pMP::MP-GFP* (**A**, **C**, **E**, **G**) and *pMP::MP11ir-GFP* (**B**, **D**, **F**, **H**) transgenic lines. The GFP fluorescence is shown in green. The blue fluorescence marks the cell (Renaissance staining). (**A-H 1-4**) Magnification views of the areas marked in A-H. Size bars indicate 200 μm (**A-F**), 20 μm (**A-H1-4**).

### Overexpression of a truncated MP inhibits the SE process

To better understand the complex roles of full-length and truncated MP proteins in SE, we examined the effects of ectopic expression of the truncated MP protein lacking the PB1 domain (ΔARF5), as MP11ir. We utilized a β-estradiol inducible *pXVE::*Δ*ARF5 pDR5::GFP* transgenic line to overexpress ΔARF5. We observed that overexpression of Δ*ARF5* inhibits the SE process when explants are cultured on a 2,4-D supplemented medium (Fig. 4). Specifically, explants overexpressing Δ*ARF5* treated with 1 µM β-estradiol and different concentrations of 2,4-D (1.0, 2.5, 5.0 µM 2,4-D) developed callus tissue rather than somatic embryos. This finding underscores that constitutive overexpression of Δ*ARF5* affects morphogenesis and disrupts the transition to embryonic development.

**Figure 4.**
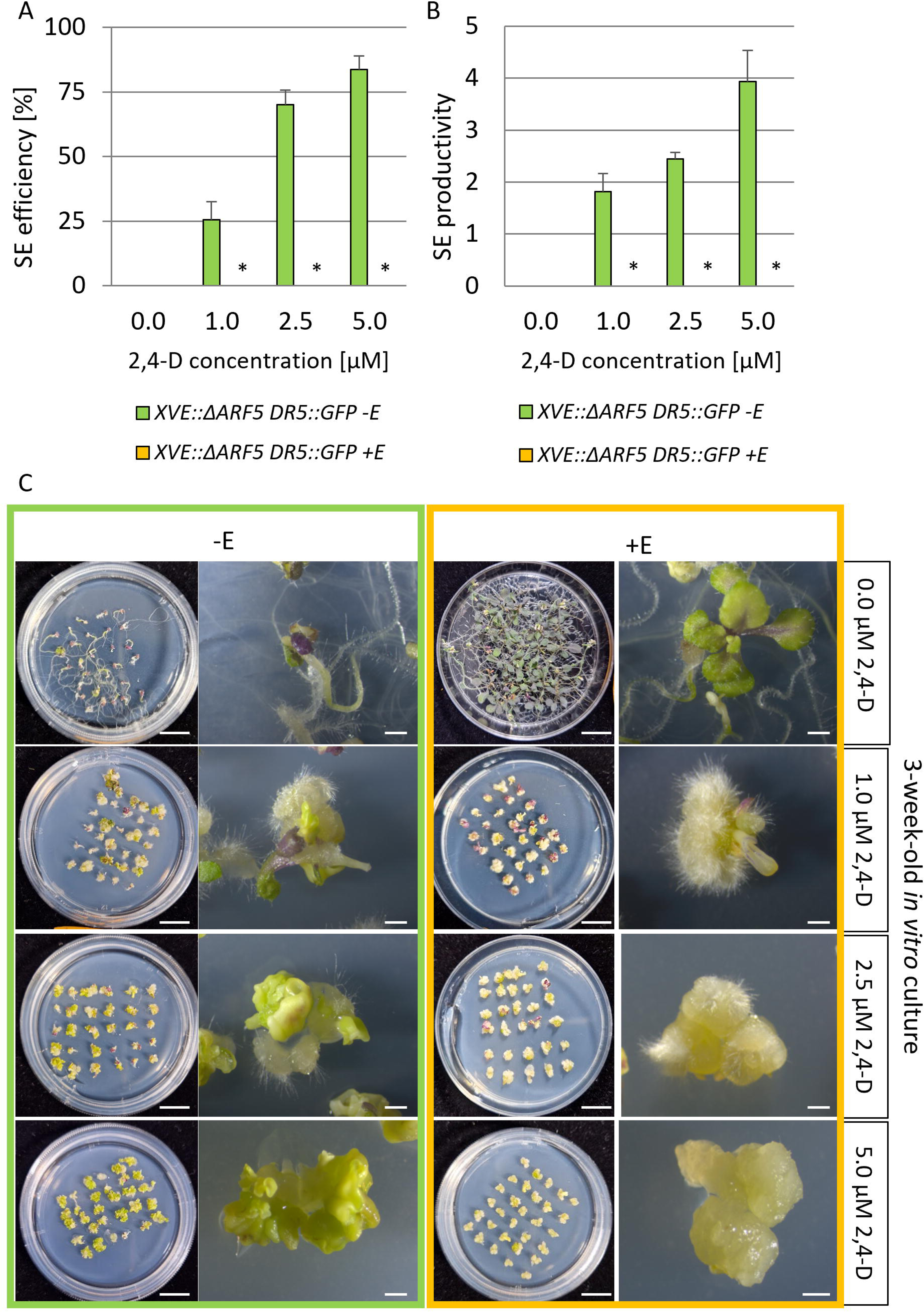
Overexpressing Δ*ARF5* impairs SE induction. (**A, B**) The embryogenic capacity of *pXVE::*Δ*ARF5 pDR5::GFP* explants treated with different 2,4-D concentrations was evaluated in a 21-day-old culture. The significantly impaired SE efficiency (**A**) and productivity (**B**) of the Δ*ARF5*-expressing explants are presented. (**C**) Explants overexpressing Δ*ARF5* and cultured on auxin-media could not regenerate somatic embryos and produced callus. Δ*ARF5* overexpression was induced with 1 µM estradiol (+E). Values significantly different from the control culture (Col-0) are marked with asterisks. A two-way ANOVA analysis (*p* < 0.05) followed by Tukey’s HSD was used to determine any values that were significantly different (\**p* < 0.05) (n = 3; means ± SD are given). Size bars indicate 1 mm for explant images and 1 cm for plate images.

### Truncated and canonical MP proteins influence genes involved in auxin biosynthesis

The inhibitory effect of Δ*ARF5* overexpression on the SE process prompted us to investigate its potential influence on endogenous auxin levels. SE inhibition often correlates with suboptimal auxin concentrations during SE induction with IZE. Indeed, overexpression of the *LEAFY COTELYDON 2* (*LEC2*) gene, a master regulator of the embryogenic process during auxin-dependent SE induction, resulted in SE inhibition due to increased endogenous auxin levels (Wójcikowska and Gaj, 2015; Wójcikowska et al., 2013). Similarly, the combined treatment of exogenous auxin and TSA inhibits SE in IZEs (Wójcikowska et al., 2018). To explore the link between ΔARF5-induced phenotypes and fluctuations in endogenous auxin levels, we examined the expression of auxin biosynthetic genes from the IPyA pathway: *TAA1, TAR1, TAR2*, and the flavin monooxygenases *YUC1-11*. These genes are expressed during the SE process and are critical for auxin production during SE formation (Bai et al., 2013; Karami et al., 2023; Li et al., 2022). Using the *pXVE::*Δ*ARF5 pDR5::GFP* transgenic line, we induced Δ*ARF5* overexpression in explants cultured on 2,4-D supplemented medium with 1 µM β-estradiol for 1, 5, and 10 days. Our analysis revealed that Δ*ARF5* overexpression significantly altered the expression of *TAA1*, *TAR1*, *YUC3*, *YUC5*, and *YUC8* genes (Fig. 5A-C). Specifically, *TAA1*, *TAR1*, *YUC5*, and *YUC8* showed increased expression levels after 24 hours of Δ*ARF5* overexpression (Fig. 5A), whereas *YUC3* and *YUC8* were upregulated on the 5^th^ day (Fig. 5B). Ten days of overexpression of Δ*ARF5* maintained the up-regulation of *TAA1* and *YUC5* genes but resulted in the down-regulation of *YUC3* (Fig. 5C). Further analysis in the *p35S::bdl-GR* line and the *mp* mutant confirmed that *TAA1*, *TAR1*, *YUC3, YUC5* and *YUC8* are strongly down-regulated under conditions of reduced MP function (Fig. 5D-F). The expression profiles of the other auxin biosynthetic genes in *pXVE::*Δ*ARF5 pDR5::GFP* (Fig. S3), *p35S::bdl-GR,* and *mp* lines (data not presented) were mainly characterized by constant or reduced expression. To understand the biological significance of MP11ir and MP in the *in vivo* regulation of *TAA1*, *TAR1*, *YUC3, YUC5*, and *YUC8* genes, their expression was additionally analyzed in 5- and 10-day-old explants of *pMP::MP11ir*, *pMP::MP,* and *pMP::MP11ir pMP::MP* lines in the *mpS319* background (Fig. 5G, H). In 5-day-old explants, the expression of all tested genes in the presence of single or combined MP and MP11ir proteins was similar to that in Col-0. In the 10-day-old explants, the expression of *TAA1*, *YUC3*, and *YUC5* returned to the level observed in the wild-type genotype Col-0 in the presence of MP11ir, in contrast to *TAR1* and *YUC8*, which remained lower than in Col-0. In the presence of MP alone and in combination with MP11ir, the analyzed genes had the same (*TAA1, YUC8*) or slightly higher (*YUC3*, *YUC5*) expression than in Col-0 and, notably, all genes were characterized by a significantly higher expression than in the *arf5* mutant. Interestingly, a synergistic effect of the MP11ir and MP on the expression of *YUC3* and *YUC8* was significant compared to their individual effect.

**Figure 5.**
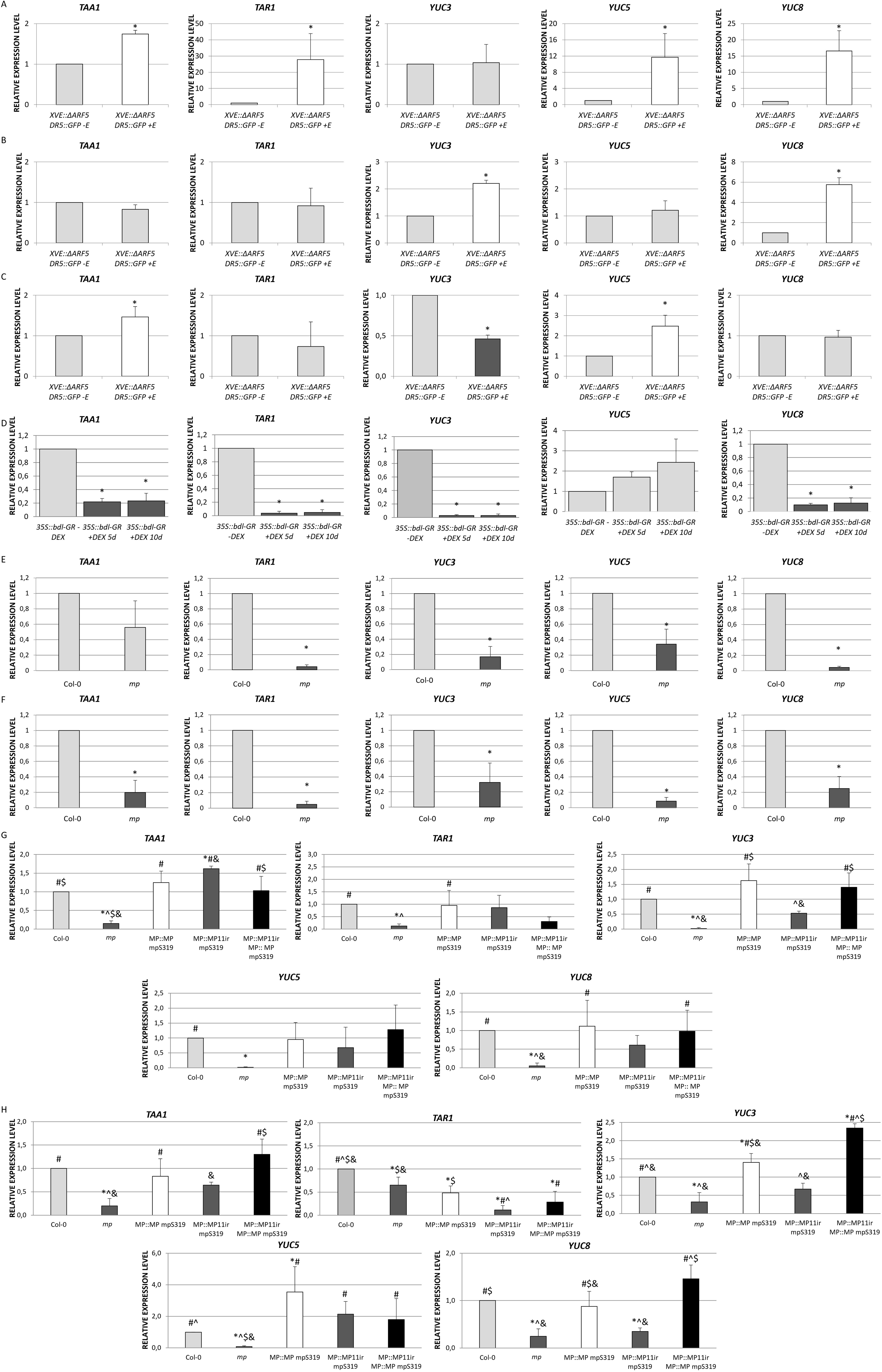
*TAA1*, *TAR1*, *YUC3*, *YUC5,* and *YUC8* expression levels are altered when MP-dependent transcriptional activities are impaired. (**A-F**) Relative expression level of *TAA1*, *TAR1*, *YUC3*, *YUC5,* and *YUC8* auxin biosynthetic genes in the 1-(**A**), 5-(**B**), and 10-day-old (**C**) embryogenic cultures of the *pXVE::*Δ*ARF5 pDR5::GFP* transgenic line treated with 1 µM β-estradiol (+E), 5- and 10-day-old embryogenic cultures of the *p35S::bdl-GR* transgenic line treated with 10 µM dexamethasone (+DEX) (**D**), and 0-(**E**) and 10-day-old (**F**) embryogenic cultures of *mp* mutant, (**G, H**). Relative expression level of *TAA1*, *TAR1*, *YUC3*, *YUC5,* and *YUC8* auxin biosynthetic genes in 5-(**G**) and 10-day-old (**H)** embryogenic cultures of *mp* mutant, *pMP::MP mpS319*, *pMP::MP11ir mpS319*, and *pMP::MP11ir pMP::MP mpS319*. The relative expression level was normalized to internal control (At4g27090) and calibrated to the *pXVE::*Δ*ARF5 pDR5::GFP* culture of the same age and untreated with 1 µM β-estradiol (-E) (**A-C**) or *p35S::bdl-GR* culture of the same age and untreated with 10 µM dexamethasone (-DEX) (**D**), or Col-0 embryogenic culture of the same age (**E, F, G**). Values significantly different from the *pXVE::*Δ*ARF5 pDR5::GFP* (-E) (**A-C**), *p35S::bdl-GR* (-DEX) (**D**) or Col-0 (**E, F**) culture of the same age are marked with asterisks (\**p* < 0.05; n = 3 ± SD). Values significantly different from the Col-0 culture of the same age are marked with asterisks (*), *arf5* are marked with hashtags (#), *pMP::MP mpS319* are marked with carets (^), *pMP::MP11ir mpS319* are marked with dollar sign ($), *pMP::MP11ir pMP::MP mpS319* are marked with ampersands (&) (**G**). (*#^$& *p* < 0.05; n = 3 ± SD).

Notably, a DAP-seq analysis (O’Malley et al., 2016) identified binding peaks for MP on the promoters of *TAA1*, *TAR1*, and *YUC5* and for ARF2 on the *YUC8* promoter (Fig. S4). All the genes deregulated in response to ΔARF5 have at least one ARF-binding peak. These results collectively suggest that MP, ΔARF5, and thus presumably MP11ir, may positively regulate the expression of genes involved in auxin biosynthesis, specifically *TAA1*, *TAR1*, *YUC3*, *YUC5,* and *YUC8* genes.

### The localization of TAA1, TAR1, YUC3, YUC5, and YUC8 enzymes during induction of the SE process is regulated by MP and MP11ir

To elucidate the spatial distribution of the enzymes involved in the IPyA-dependent auxin biosynthesis during the embryogenic transition, we utilized translational reporter lines for TAA1, TAR1, TAR2, and YUCs proteins (Fig. S5). The TAA1, TAR1, YUC3, YUC5, and YUC8 enzymes were selected based on the RT-qPCR results (Fig. 5). To validate the regulation of their expression by MP and ΔARF5, lacking the PB1 domain similar to MP11ir, we crossed the translational reporter lines for TAA1, TAR1, YUC3, YUC5, and YUC8 into *pXVE::*Δ*ARF5 pDR5::GFP, p35S::bdl-GR*, and *mp* lines to investigate any changes in the localization of GUS signals. Explants were induced into SE with 2,4-D and 1 µM β-estradiol or 10 µM DEX to induce Δ*ARF5* or *bdl* overexpression, respectively, and sampled on the 5^th^ and 10^th^ days of the culture. Additionally, *mp* IZE (0 day) and explants on the 5^th^ and 10^th^ days of the SE process were analyzed.

Among the enzymes analyzed, TAA1, YUC3, YUC5, and YUC8 exhibited distinct expression patterns in response to Δ*ARF5* overexpression (Fig. 6). Notably, YUC8 showed robust accumulation in the proximal regions of cotyledons and hypocotyl upon Δ*ARF5* overexpression, as evidenced by intense GUS signals (Fig. 6A, Fig. 7E). Conversely, *bdl* overexpression led to a significant decrease in GUS signal intensity for YUC8 (Fig. 6B, Fig. 7E). In *mp* mutant, the GUS signal for YUC8 was absent (Fig. 6C, Fig. 7E), confirming the MP*-* and MP11ir-dependent expression of the *YUC8* gene. TAA1 was broadly active throughout the IZE explants during embryogenic transition (5^th^ and 10^th^ day), with a slight increase of the GUS signal upon Δ*ARF5* overexpression (Fig. 6A, Fig. 7A). However, changes in GUS intensity were modest upon *bdl* overexpression or in *mp* mutant (Fig. 6B, C, Fig. 7A). YUC3 exhibits ubiquitous localization in the hypocotyl region of explants, which is enhanced in the cotyledons upon Δ*ARF5* overexpression (Fig. 6A, Fig. 7C). In contrast, *bdl* overexpression deeply reduced the intensity of the GUS signal (Fig. 6B, Fig. 7C), which was completely absent in *mp* mutant IZEs or reduced at 5^th^ and 10^th^ days of SE induction (Fig. 6C, Fig. 7C). YUC5 showed a specific accumulation at the tip of the cotyledons only on the 10^th^ day-old SE under Δ*ARF5* overexpression (Fig. 6A, Fig. 7D), with no changes observed during *bdl* overexpression (Fig. 6B, Fig. 7D). In *mp* mutant, the GUS signal for YUC5 was restricted to single cells (Fig. 6C, Fig. 7D). In contrast, TAR1 localization was unchanged upon Δ*ARF5,* and *bdl* overexpression, and in *mp* mutant (Fig. 6, Fig. 7B).

**Figure 6.**
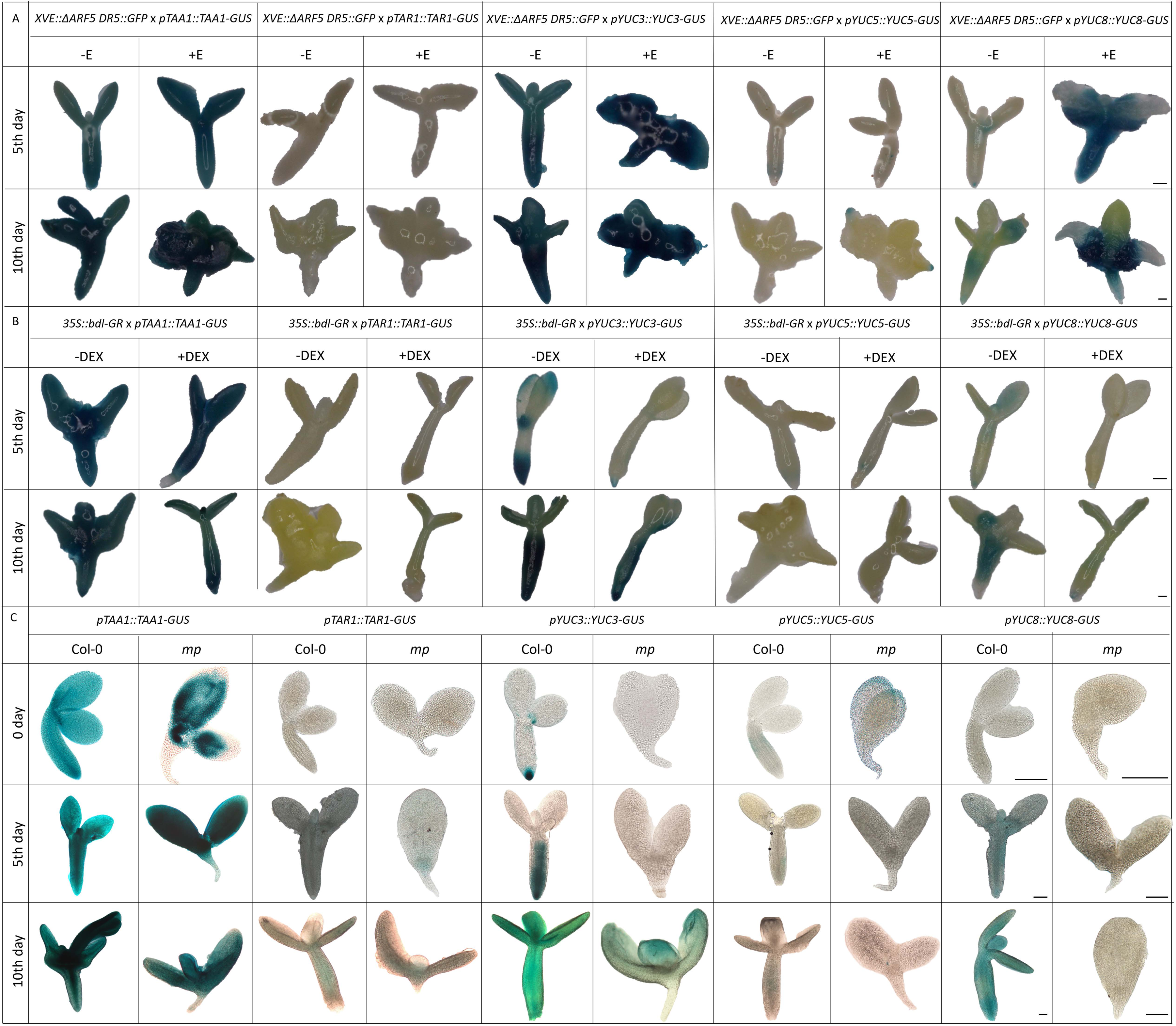
*TAA1*, *TAR1*, *YUC3*, *YUC5*, and *YUC8* expression patterns are altered when MP-dependent transcriptional activities are impaired. Spatio-temporal localization of TAA1, TAR1, YUC3, YUC5, and YUC8 enzymes during SE process under *in vitro* culture in *pXVE::*Δ*ARF5 pDR5::GFP* (**A**) *p35S::bdl-GR* (**B**) or *mp* background (**C**). IZE explants of reporter lines were cultured on an E5 medium as a control (**A, B, C**), E5 medium with 1 µM β-estradiol for Δ*ARF5* overexpression (+E) (**A**), E5 medium with 10 µM dexamethasone (+DEX) (**B**). The tissue was sampled on the 5- and 10-day-old *in vitro* cultures (**A, B**), and in IZEs, explants on the 5- and 10-day-old *in vitro* cultures (**C**). The expression is visible in blue from *pTAA1::TAA1-GUS*, *pTAR1::TAR1-GUS*, *pYUC3::YUC3-GUS*, *pYUC5::YUC5-GUS*, and *pYUC8::YUC8-GUS* reporter lines crossed into the respective genetic backgrounds. Size bars indicate 200 μm.

**Figure 7.**
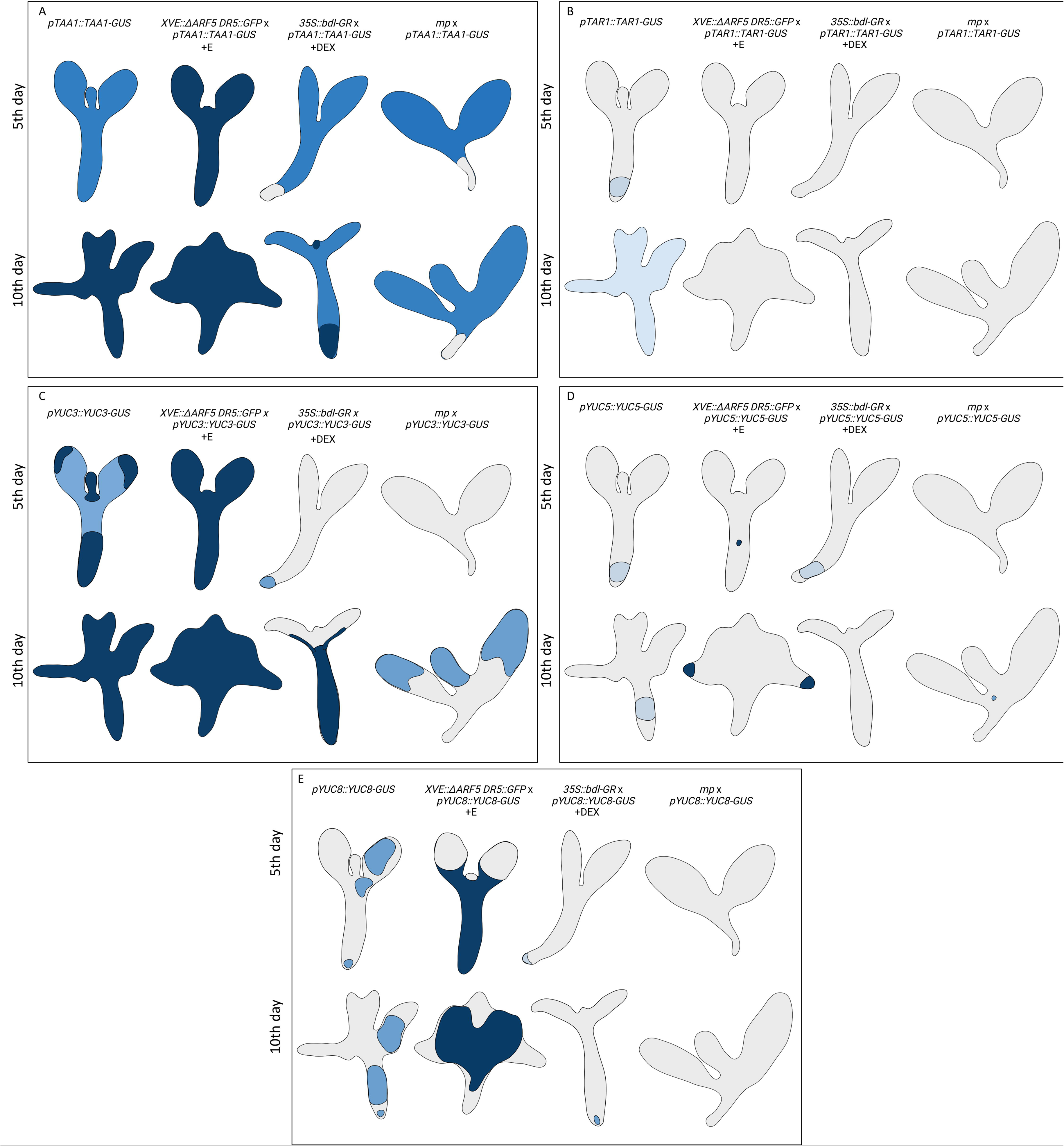
Summary of the localization of TAA1 (**A**), TAR1 (**B**), YUC3 (**C**), YUC5 (**D**), and YUC8 (**E**) enzymes during SE process under *in vitro* culture of Col-0, *pXVE::*Δ*ARF5 pDR5::GFP* line treated with β-estradiol, *p35S::bdl-GR* line treated with DEX, and *mp* mutant.

These findings demonstrated the cell-type-specific and temporal regulation of the TAA1, YUC3, YUC5, and YUC8 auxin biosynthetic enzymes by ΔARF5, MP, and MP11ir proteins during the SE process.

### Influence of MP11ir on endogenous auxin levels

We hypothesized that MP11ir might regulate the endogenous auxin levels through TAA1, YUC3, YUC5, and YUC8 during SE. We explored this hypothesis using three approaches. First, we assessed local auxin responses in the *pXVE::*Δ*ARF5 pDR5::GFP* explants during the auxin-independent embryogenic transition (1^st^ and 5^th^ days of TSA treatment) to avoid activation of the *DR5* signal by the exogenously applied auxin. Explants were cultured on an ET medium with or without β-estradiol. GFP signals indicate auxin-responsive gene transcription in the root and shoot apical meristems and the tip of cotyledons in non-induced explants (Fig. 8A). After 24 hours of induced Δ*ARF5* overexpression, an enhanced GFP signal was observed in the hypocotyl and throughout the cotyledons (Fig. 8B, E). In 5-day-old explants, Δ*ARF5* overexpression intensified the GFP signal in the hypocotyl and cotyledons (Fig. 8C, D, F). Second, we quantified indolic compounds and IAA in 21- or 10-day-old *in vitro* cultures, respectively, of *pXVE::*Δ*ARF5 pDR5::GFP* explants treated and not with β-estradiol (Fig. 8G, H). The explants overexpressing Δ*ARF5*, exhibiting callus formation, showed a 1.37-fold increase in indolic compound levels and a 3-fold increase in IAA content compared to non-induced explants. Finally, to determine whether the formation of calli might result from high auxin responses and indolic compound levels, we analyzed the SE process in Col-0 and the *pXVE::*Δ*ARF5 pDR5::GFP* line treated with different concentrations of yucasin (100 and 150 µM), an inhibitor of YUC activity (Tsugafune et al., 2017). (Non-)induced Col-0 (-E, +E) and non-induced *pXVE::*Δ*ARF5 pDR5::GFP* (-E) explants treated with yucasin showed highly reduced SE efficiency (1.3 to 1.5 times lower) and productivity (no more than two somatic embryos per reacting explant were formed) (Fig. S6). The results align with published literature (Karami et al., 2023; Li et al., 2022). We then investigated whether treating *pXVE::*Δ*ARF5 pDR5::GFP* IZE explants with yucasin could reverse the phenotype induced by Δ*ARF5* overexpression with β-estradiol and enable the explants to regenerate embryos instead of callus tissue. Indeed, these *pXVE::*Δ*ARF5 pDR5::GFP* IZE explants formed a few somatic embryos (15.4% with 100 and 3.4% with 150 µM yucasin) and with low productivity (1 somatic embryo per reacting explant) instead of the calli observed in the absence of yucasin (Fig. S6).

**Figure 8.**
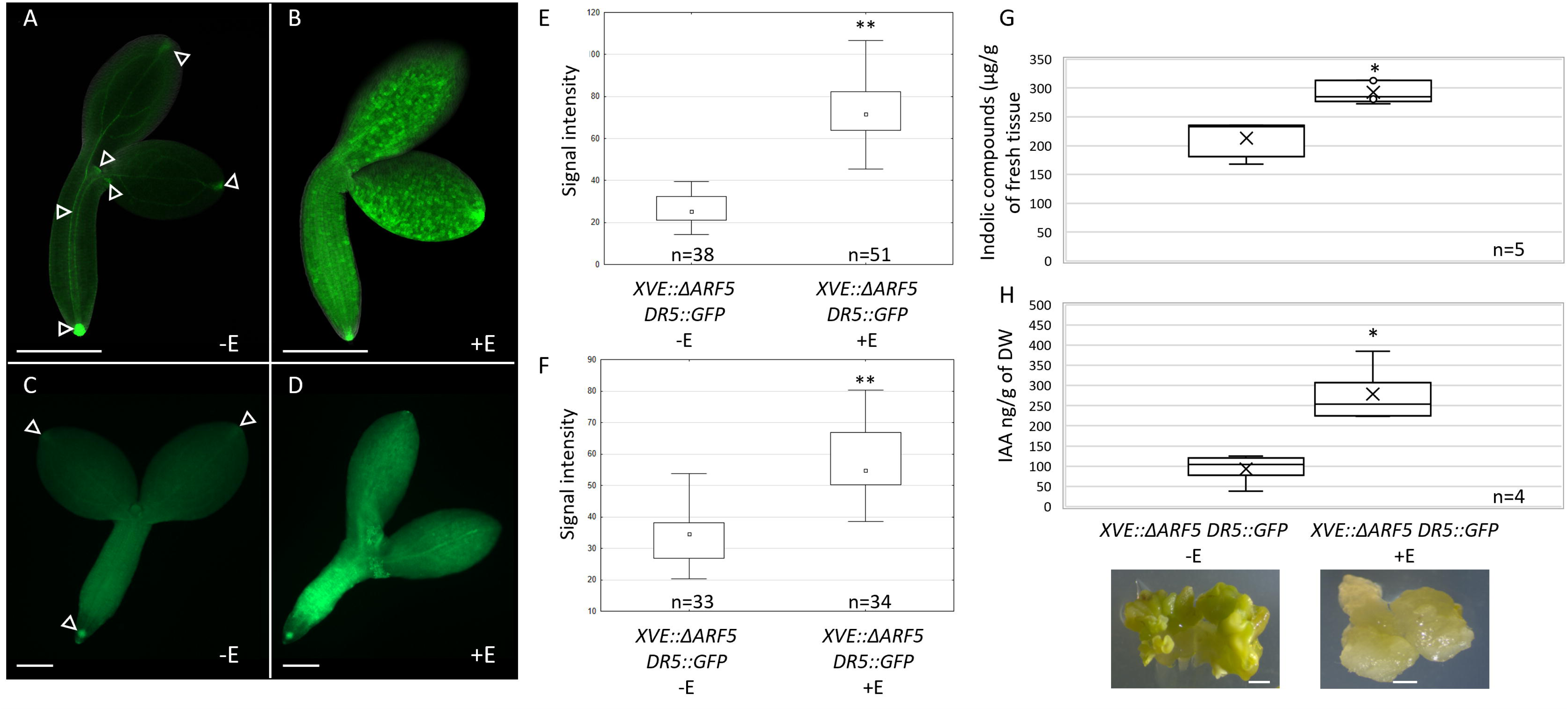
Overexpressing Δ*ARF5* increases endogenous auxin levels. (**A-D**) GFP-monitored auxin maxima in *pXVE::*Δ*ARF5 pDR5::GFP* explants cultured for 1 (**A, B**) and 5 days (**C, D**) on an ET medium (1.0 μM TSA) without (-E) (**A, C**) or with β-estradiol (+E) (**B, D**). The GFP fluorescence signal is displayed in green. (**E, F**) The fluorescence signal was quantified in 1-(**E**) and 5-day-old (**F**) explants. (**G**) Level of indolic compounds (μg/g of fresh tissue) in 3-week-old IZE-culture of *pXVE::*Δ*ARF5 pDR5::GFP* cultures with 1 µM β-estradiol to induce Δ*ARF5* overexpression. (**H**) Level of IAA (ng/g of dry weight DW) in 10-day-old IZE-culture of *pXVE::*Δ*ARF5 pDR5::GFP* with 1 µM β-estradiol to induce Δ*ARF5* overexpression. A two-way ANOVA analysis (*p* < 0.05) followed by Tukey’s HSD was used to determine any values that were significantly different (\**p* < 0.05; \*\**p* < 0.01). Size bars indicate 200 μm (**A-D**) and 1 mm (**G**). Arrowhead (**A, C**) marked regions of high auxin response in *pXVE::*Δ*ARF5 pDR5::GFP* explants untreated by β-estradiol (-E).

This finding supports that Δ*ARF5* overexpression amplifies local auxin responses by modulating auxin production and signaling during SE, ultimately forming calli instead of embryonic structures. However, other genetic components deregulated by Δ*ARF5* overexpression also likely inhibit SE capacity.

## Discussion

*MP* is upregulated in plant embryogenic cells and serves as a potential molecular marker for this developmental stage. Various transcriptomic studies support the involvement of *MP* in the SE process in various plant species (Awada et al., 2023; Capote et al., 2019; Chen et al., 2020; Gliwicka et al., 2013; Indoliya et al., 2016; Juárez-González et al., 2019; Qi et al., 2021; Quintana-Escobar et al., 2019; Shi et al., 2016; Thibaud-Nissen et al., 2003; Wickramasuriya and Dunwell, 2015; Xu et al., 2024; Zeng et al., 2024; Zhu et al., 2023, Ge et al., 2025). Additionally, the presence of *MP* transcripts in regions critical for SE induction, such as cotyledons and SAM, suggests that MP may be involved in the embryogenic transition (Wójcikowska and Gaj, 2017). Two *mp* mutant alleles, *arf5* (SALK_001058C) (Wójcikowska and Gaj, 2017) and *mpS319* (SALK_021319) (present work), exhibit impaired SE processes. This inhibition is linked to the differential expression of SE master regulators (Wójcikowska and Gaj, 2017). Similarly, the *Osarf5* mutant displays partial defects in forming a scutellum-derived embryogenic callus. The *Osarf5* mutant also shows a significantly reduced expression of *LEAFY COTELYDON 1* (*OsLEC1*), indicating that the OsARF5-mediated auxin signaling pathway might regulate *OsLEC1* (Guo et al., 2023a).

Expressing auxin-insensitive bdl protein during SE induction revealed that SE inhibition in the *mp* mutant is not due to its embryonic defects but is triggered by MP inactivity in IZE. Many *EMBRYO-DEFECTIVE* (*EMB*) genes involved in zygotic embryo development and morphogenesis, such as *LEC1*, *LEC2*, *FUSCA 3*, *DICER-LIKE 1*, *WUSCHEL RELATED HOMEOBOX 5* (*WOX5*), and *PIN-FORMED 1* (*PIN1*), are also critical for SE processes (Ledwoń and Gaj, 2009; Ledwoń and Gaj, 2011; Meinke, 2020; Su et al., 2015; Su et al., 2009; Szyrajew et al., 2017). However, the overexpression of the auxin-insensitive bdl protein had a more substantial impact on inhibiting embryogenic transition than the *mp* mutation, suggesting that mutated bdl protein might repress other ARFs besides MP (Piya et al., 2014).

MP’s transcriptional targets likely include genes encoding proteins essential to the SE process. AGAMOUS-LIKE 15 (AGL15) and AGL18, crucial for SE (Harding et al., 2003; Paul et al., 2022; Thakare et al., 2008; Zheng and Perry, 2014), might regulate *MP* expression as the *agl15 agl18* double mutant shows a significant reduction of *MP* mRNA levels (Zheng et al. 2009). Also, inhibition of PHYTOCHROME-INTERACTING FACTOR 4 (PIF4) induces several auxin responses (*ARF5, ARF8,* and *ARF16*) and biosynthetic (*YUC1, YUC2*, *YUC6*, *CYP79B2,* and *AMIDASE 1* (*AMI1*)) genes, resulting in the formation of the somatic embryos (Mira et al., 2023). Despite these insights, the detailed mechanism of action of MP and its MP11ir isoform during SE induction and somatic embryo formation remains unknown. Lines expressing truncated MP proteins without the PB1 domain could elucidate the complex action of MP11ir by mimicking its function. Truncated ΔARF5 and MP^abn^ proteins partially rescue *mp* phenotypic defects but cause unique leaf defects (Garrett et al., 2012; Krogan et al., 2012). Ectopic ΔARF5 protein stimulates the formation of new shoots *in vivo* (Ma et al., 2019), whereas *mpS319* mutants show reduced shoot regeneration frequency (Zhang et al., 2021). During *in vitro* culture, ΔARF5 increases the shoot organogenesis frequencies in Arabidopsis root-derived explants and activates auxin signaling. Visualization and cytometric quantification of the *pDR5::GFP* nuclear auxin signaling reporter revealed an increased GFP signal in roots and protoplasts overexpressing Δ*ARF5* (Gonzalez et al., 2021). A similar trend was observed after inducing Δ*ARF5* expression during the embryogenic transition. MP is involved in shoot apical meristem (SAM) activity and maintenance by directly controlling the expression of ARABIDOPSIS HISTIDINE PHOSPHOTRANSFER PROTEIN 6 (AHP6) (Besnard et al., 2014), TARGET OF MONOPTEROS (TMOs) (Schlereth et al., 2010), DORNRÖSCHEN/ENHANCER OF SHOOT REGENERATION 1 (DRN/ESR1) (Cole et al., 2009), *WUSCHEL RELATED HOMEOBOXs* (*WOXs*) (Lee et al., 2022; Tvorogova et al., 2019), essential for shoot regeneration or SE (Fambrini et al., 2022). The shoot regeneration-promoting effect of ΔARF5 (Ckurshumova and Berleth, 2015; Gonzalez et al., 2021) may help to overcome organogenic resistance in recalcitrant explants and species in the future. However, Δ*ARF5* overexpression did not enhance embryogenic capacity (our results) but instead led to callus formation, similar to *NcARF5* overexpression in *Neolamarckia cadamba*, which triggers rapid callus proliferation (Ma et al., 2024).

Studies on embryo-derived SE systems showed that SEs initiate from the callus edge, where auxin accumulates, as visualized using PIN1 and DR5 reporters (Kadokura et al., 2018; Su et al., 2015; Su et al., 2009). MP directly and positively regulates PIN1, which is responsible for auxin efflux in polar auxin transport (PAT) (Krogan et al., 2016). Auxin is present in the *pin1* mutant meristem, ensured by vascular tissue transport or induction of auxin biosynthesis (Banasiak et al., 2019). Therefore, MP is proposed as a primary regulator of auxin transport and biosynthesis. Our study identifies auxin biosynthetic genes, *TAA1*, *TAR1*, *YUC3*, *YUC5,* and *YUC8*, as potentially positively regulated by ΔARF5, MP11ir, and/or MP during *in vitro* embryogenic transition. We propose that MP and MP11ir proteins directly or indirectly control the TAA1/TARs-YUCs-mediated auxin biosynthetic pathway (our work) and polar auxin transport during embryogenesis (Robert et al., 2015; Schlereth et al., 2010). This hypothesis aligns with findings that MP affects *YUC1* and *YUC8* expression to specify the ground tissue in early globular zygotic embryos (Möller et al., 2017). IPyA-dependent auxin biosynthesis is crucial for effective SE induction and somatic embryo identity and growth (Li et al., 2022). Yucasin treatment, inhibiting YUC activity (Tsugafune et al., 2017), reduces SE efficiency and productivity in Arabidopsis (Karami et al., 2023; Li et al., 2022) and coffee culture (Uc-Chuc et al., 2020). The importance of auxin biosynthesis in the SE process is evidenced by the decreased embryogenic response in *yuc* mutants. Multiple *yuc* mutants, *yuc3 yuc8* (Li et al., 2022), *yuc1 yuc4 yuc10 yuc11* (Bai et al., 2013), *yuc3 yuc5 yuc7 yuc8 yuc9* (Sakamoto et al., 2022), display disrupted SE or callus formation compared to single mutants (Sakamoto et al., 2022; Wójcikowska et al., 2013). *YUC* genes are direct targets of TFs, such as LEC1, LEC2, BABY BOOM (BBM), and AT-HOOK MOTIF CONTAINING NUCLEAR LOCALIZED 15 (AHL15), which play pivotal roles in embryogenic transition, demonstrated by decreased SE in their mutant and spontaneous induction of somatic embryo formation when overexpressed (Boutilier et al., 2002; Gaj et al., 2005; Karami et al., 2021; Ledwoń and Gaj, 2009; Lotan et al., 1998; Stone et al., 2001). During SE induction, LEC1 activates *YUC4* and *YUC10* (Hu et al., 2018; Junker et al., 2012), while LEC2 positively regulates *YUC1*, *YUC4*, and *YUC10* expression (Stone et al., 2008; Wójcikowska et al., 2013). BBM induces local and ectopic expression of *YUC3* and *YUC8* coupled with IAA biosynthesis and activation of the WOX2 embryo marker (Li et al., 2022). During SE induction, AHL15 activates *YUC6*, *YUC7*, *YUC8*, and *YUC9* expression (Karami et al., 2023), confirming YUCs as crucial molecular elements of a network governing the SE process.

We showed that both *MP11ir* and *MP* transcript levels increase along with the concentration of auxins (2,4-D, NAA, IAA). The auxins analyzed in our study exhibited different potencies in inducing SE, and with increasing auxin concentrations, a greater ability to induce SE was observed. Similarly, the synthetic auxin 2,4-D was effective in SE induction, whereas natural auxins were less efficient or even failed to induce regeneration (Karami et al., 2024). Why do auxins act differently? 2,4-D cannot be efficiently transported and accumulates in the cells, triggering a strong (optimal for SE) auxin response. In contrast, natural auxins are subject to polar transport, which prevents their local accumulation and results in a moderate auxin response - insufficient for SE induction. It has been shown that both natural auxins and synthetic analogs become efficient inducers of SE when their efflux is transiently inhibited by an auxin transport inhibitor. Explants of auxin efflux mutants, such as *pin2* and *abcb1 abcb19*, show enhanced SE efficiency when treated with IAA or efflux-inhibited IAA, confirming that auxin efflux reduces the efficiency of SE in Arabidopsis. Efflux-inhibited IAA, similar to 2,4-D, also efficiently induced SE from carrot suspension cells, and efflux-inhibited 4-Cl-IAA improved *de novo* shoot regeneration in *Brassica napus* (Karami et al., 2024).

A study identified target genes of MP and truncated ΔARF5 proteins in 3-day-old seedlings (un)treated with auxin (Xie et al., 2022). This ChIP-seq analysis suggests that deleting the PB1 domain renders the protein auxin-insensitive and alters its binding profile. Among genes involved in the IPyA auxin biosynthesis pathway, *YUC3* and *YUC5* promoters are recognized and bound by ΔARF5, while the *YUC6* promoter is targeted by ΔARF5 and MP. Differences in the preferential AuxRE motifs bound by ΔARF5 and MP suggest that the PB1 domain may affect homodimer formation and DNA binding, indicating that ΔARF5 and MP may differ in their molecular actions and thus impact plant development differently (Xie et al., 2022).

Similarly, recent transcriptomic data revealed that MP and MP11ir may regulate the expression of different sets of genes during Arabidopsis root development and have specific and non-redundant functions. Among DEGs, *TAR2* (1.6 and 1.7-fold) and *YUC9* (1.5- and 1.8-fold) can be distinguished with an increased expression in *pMP::MP* or *pMP::MP11ir* roots, respectively, relative to the wild type. Other genes involved in Trp (IAA precursor) (*IGPS*, *TRP1*, *TSA1*, *TSBtype2*) and IAA biosynthesis or conjugation (*AMI1*, *NIT1*-3, *IAMT1*, *MES17*, *CYP79B3*) are also deregulated in an MP or MP11ir-dependent manner in root cells (Cavalleri et al., 2024). It suggests that MP and MP11ir may regulate the expression of genes involved in auxin metabolism and modulate cell’s responsiveness to auxin.

Surprisingly, the lack of a GFP signal in the *pMP::MP11ir-GFP* SE makes it impossible to follow the isoform localization. This may be due to a weak GFP signal, below the detection threshold, or to post-transcriptional or post-translational modifications of the MP11ir-GFP proteins. The *pMP::MP:EGFP(MR)* line, in which the EGFP sequence is inserted into the MR domain of MP, upstream of the differentially spliced 11th intron, was used to localize MP11ir in root (Cavalleri et al., 2024). Such a line, together with the *pMP::MP:VENUS-2Ap-mTURQUOISE,* may allow the localization of MP and MP11ir proteins during the SE process.

Transcriptomic data have shown that many splice isoforms formed due to alternative splicing (AS) show cell-specific expression (Klepikova et al., 2016; Martín et al., 2021). Among AS events, intron retention events are functionally underestimated in plants. Given the importance of AS in responding to various stresses, it is likely required for proper environmental response in plants (Guo et al., 2023b). Limited functional evidence supports their specific expression patterns or distinct ability to rescue mutant loss-of-function phenotypes. For example, *ARF8* undergoes tissue-specific AS in flower, where the *ARF8*.*4* variant, which retains the eighth intron and has a premature stop codon, is over-represented. Overexpression of *ARF8.4*, but not other *ARF8* isoforms, reverts stamen elongation defects associated with the *arf8* knockout mutation (Ghelli et al., 2023; Ghelli et al., 2018). Another example is the *BIGPETAL* (*BPE*) gene, which encodes a petal-specific *BPEp* transcript that retains its last intron, in addition to the canonical *BPEub* transcript uniformly expressed in all organs (Szécsi et al., 2006). The C-terminal sequence exclusively encoded by *BPEp* is required to interact with ARF8. The *arf8* loss-of-function mutant phenocopies the petal defects of the *bpe* knockout mutants, suggesting that BPEp, unlike the canonical BPEub protein, is required for petal development (Kashkan et al., 2022; Varaud et al., 2011).

In conclusion, both MP proteins, canonical MP and MP11ir isoform, seem to be involved in the SE process (Fig. 9A). Potentially different expression patterns of MP and MP11ir suggest that they act in different cells, probably with different endogenous auxin levels. The ectopic expression of Δ*ARF5,* results in a global increase in endogenous auxin content in explants through direct or indirect activation of *TAA1*, *TAR1*, *YUC3*, *YUC5,* and *YUC8* genes. Disruption of auxin homeostasis in explant cells by overproduction of TAA1, YUC3, YUC5, and YUC8 enzymes leads to SE inhibition and callus formation (Fig. 9B). Blocking MP action by expressing the *bdl* repressor results in decreased levels of TAA1, YUC3, YUC5 and YUC8 enzymes during SE induction, inhibiting the SE process and promoting seedling development (Fig. 9C). The absence of both MP and MP11ir in *mp* mutant leads to the decreased TAA1 level, absence of *YUC3*, *YUC5,* and *YUC8* during early stage of SE induction, resulting in SE inhibition and callus formation (Fig. 9D). However, the analysis of auxin biosynthetic gene expression in the complemented lines, expressing MP (*pMP::MP mpS319*), MP11ir (*pMP::MP11ir mpS319*) or both proteins together (*pMP::MP pMP::MP11ir mpS319*), suggest that MP and MP11ir may have common and specific transcriptional targets during SE, with MP11ir regulating *TAA1*, *YUC3*, and *YUC5*, MP regulating *YUC8* and a synergistic MP and MP11ir effect on *YUC3* and *YUC5*. In conclusion, by suggesting that MP and MP11ir are central regulators of auxin biosynthesis during embryogenic processes, this study provides an elegant and straightforward experimental setup that elucidates the role of MP and MP11ir in SE induction, acting through the transcriptional control of auxin biosynthetic genes.

**Figure 9.**
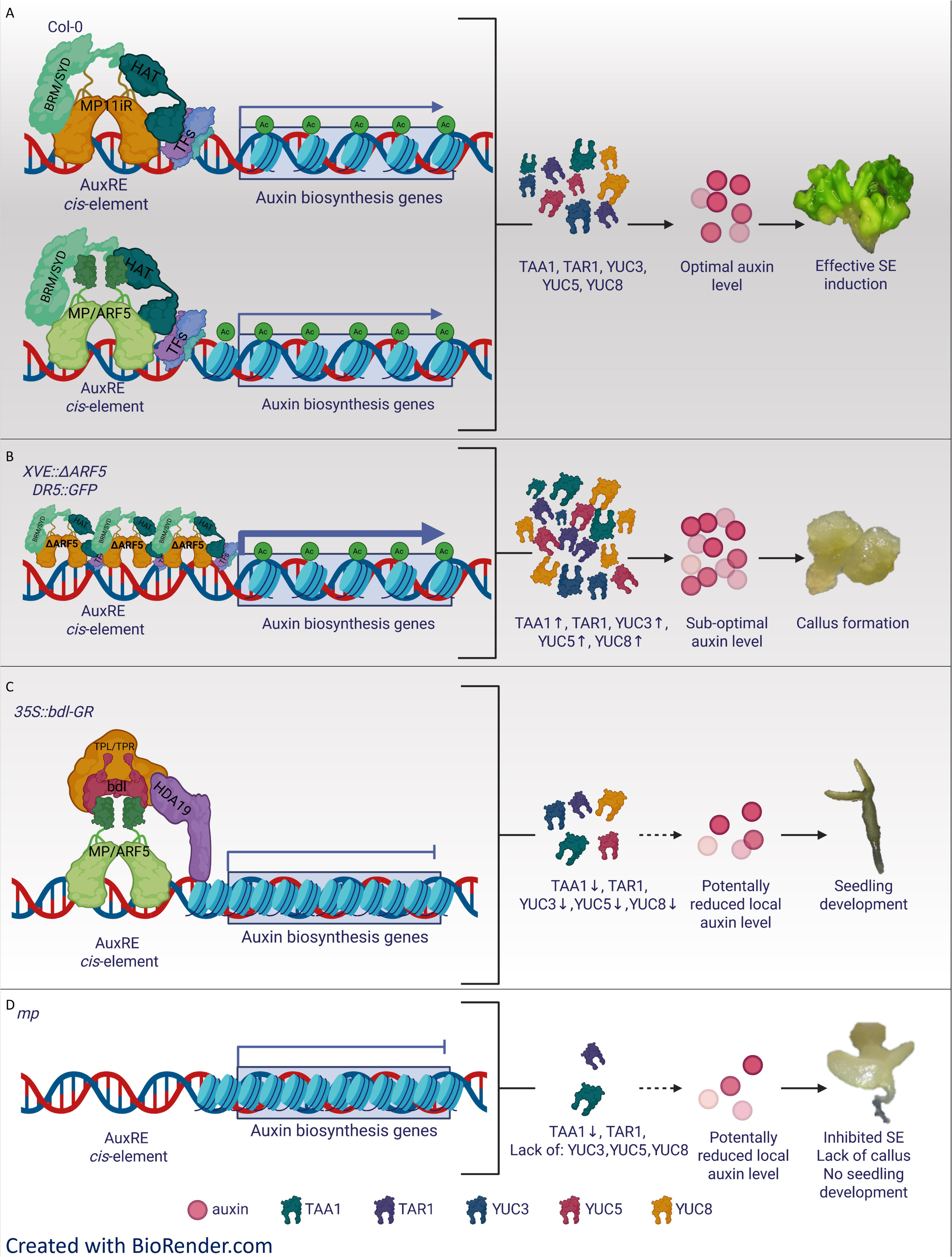
Proposed model of MP and MP11ir action during embryogenic transition through control of auxin-biosynthesis genes expression. (**A**) The canonical MP protein and MP11ir isoform are required for the efficient embryogenic transition by controlling auxin biosynthesis genes. (**B**) Overexpression of a truncated Δ*ARF5* protein, caused elevated level of TAA1, YUC3, YUC5, and YUC8 enzymes in explants, which disrupts auxin homeostasis, leads to suboptimal endogenous auxin levels, inhibits the SE process, and induces the formation of callus. (**C**) Blocking MP action through the expression of the *bdl* repressor results in decreased TAA1, YUC3, YUC5 and YUC8 levels during SE induction, inhibits the SE process, and induces seedling development. (**D**) The lack of MP and MP11ir in *mp* mutant leads to the decreased TAA1 level, absence of YUC3, YUC5, and YUC8 enzymes during early stage of SE induction what inhibits the SE process, and leads to callus formation. AuxRE – Auxin Response Elements; BRM/SYD – BRAHMA/SPLAYED; MP – MONOPTEROS; MP11ir – MP isoform; TFs - TRANSCRIPTION FACTOR; HAT – HISTONE ACETYLASES; TAA1 - TRYPTOPHAN AMINOTRANSFERASE OF ARABIDOPSIS 1, TAR1 - TRYPTOPHAN AMINOTRANSFERASE RELATED 1; TPL/TPR – TOPLESS/TPL-RELATED; YUC – YUCCA; Ac – acetyl group; SE – somatic embryogenesis. Created with BioRender.com.

## Materials and methods

### Plant material

The seeds of *Arabidopsis thaliana* (L.) Heynh Columbia (Col-0) wild type, *mpS319* mutant (N21319), reporter lines *pTAA1::TAA1-GUS* (N72246), *pTAR1::TAR1-GUS* (N72247), *pTAR2::TAR2-GUS* (N72248), *pYUC1::YUC1-GUS* (N72250), *pYUC2::YUC2-GUS* (N72252), *pYUC3::YUC3-GUS* (N72254), *pYUC4::YUC4-GUS* (N72256), *pYUC5::YUC5-GUS* (N72259), *pYUC6::YUC6-GUS* (N72261*), pYUC7::YUC7-GUS* (N72262), *pYUC8::YUC8-GUS* (N72264), *pYUC9::YUC9-GUS* (N72266), *pYUC01::YUC10-GUS* (N72268), and *pYUC11::YUC11-GUS* (N72271) were supplied by NASC (The Nottingham Arabidopsis Stock Centre) (Alonso et al., 2003; Brumos et al., 2019). Seeds of the *pMP::MP mpS319, pMP::MP11ir-GFP mpS319*, and *pMP::MP11ir-GFP* transgenic lines were provided by Lucia Colombo (Dipartimento di BioScienze, Università degli Studi di Milano, Milano, Italy). Dolf Weijers (Laboratory of Biochemistry, Wageningen University, and Research, Wageningen, The Netherlands) generously donated seeds of the *pMP::MP-GFP*, *p35S::BDL-GR*, *p35S::bdl-GR* lines. The seeds of the *pXVE::*Δ*ARF5 pDR5::GFP* line were provided by Bastiaan Bargmann (School of Plant and Environmental Sciences, Virginia Polytechnic Institute and State University, Blacksburg, USA). The generation of *pMP::MP mpS319* x *pMP::MP11ir-GFP mpS319*, and *pXVE::*Δ*ARF5 pDR5::GFP*; *p35S::bdl-GR; arf5* x *pTAA1::TAA1-GUS*, *pTAR1::TAR1-GUS*, *pYUC3::YUC3-GUS*, *pYUC5::YUC5-GUS*, *pYUC8::YUC8-GUS* was performed by crossing.

### Plant growth and *in vitro* culture conditions

Seeds were sterilized with sodium chloride (20% commercial bleach) and plated onto ½ MS medium (Murashige and Skoog, 1962), (Duchefa Biochemie; #M0222), 10 g L^-1^ sucrose (Penta # 57-50-1) and 8 g L^-1^ agar (Duchefa Biochemie; #P1001). Seed-derived plants grown *in vitro* were a source of seedlings, and *in vivo* were a source of leaves, flowers, and siliques for IZE isolation. Plants were grown in soil with vermiculite (4:1) at 22 °C under a 16 h photoperiod of 100 µM m^-2^ s^-1^ white, fluorescent light.

### Somatic embryogenesis induction

IZEs at the cotyledonary stage of development were used as explants for the *in vitro* culture. The IZEs were cultured on an E5 medium containing 3.2 g L^-1^ of B5 micro and macro-elements (Duchefa Biochemie; #G0210) (Gamborg et al., 1968), 20 g L^-1^ sucrose, 5.0 μM of 2,4-D (2,4-dichloro phenoxy acetic acid, Sigma-Aldrich #D7299) and 8 g L^-l^ agar (Duchefa Biochemie; #P1001), pH 5.8 (Gaj, 2001). To induce the SE process without auxin, the IZEs were cultured on an ET medium containing 3.2 g L^-1^ of B5 micro and macro-elements (Duchefa Biochemie; #G0210) (Gamborg et al., 1968), 20 g L^-1^ sucrose, 1 μM of TSA (Trichostatin A, Sigma Aldrich; #T1952) and 8 g L^-l^ agar (Duchefa Biochemie; #P1001), pH 5.8 (Wójcikowska et al., 2018). Plant materials that grow in sterile conditions were kept at 23 °C under a 16 h photoperiod of 40 µM m^-2^ s^-1^ white, fluorescent light. The explant capacity for SE was evaluated in an IZE-derived culture that was induced for 21 days on an induction media, and two parameters were calculated – SE efficiency (percentage of explants that developed somatic embryos) and productivity (average number of somatic embryos produced by embryogenic explant). Thirty explants were analyzed in at least three biological replicates of the experimental combination. Around 10,000 IZEs were isolated for all the experiments. Assuming the average embryo isolation rate (100 IZEs per hour), the time to obtain the research material for each experiment was 100 hours.

### IZE treatments

IZEs at the cotyledonary stage were treated with different 2,4-D (Duchefa Biochemie; #D0911), IAA (Duchefa Biochemie; #I0901) and NAA (Duchefa Biochemie; #N0903) (0.0, 2.5, 5.0, 7.5 μM) concentrations for ten days to evaluate the level of *MP* and *MP11ir* transcripts. IZEs were cultured on a medium with 2,4-D and treated with different yucasin (Merck; #23711-26-4) concentrations (0, 100, 150 μM) for 3 weeks to evaluate the effect of YUC activity inhibition on the embryogenic capacity. Explants were first precultured for 1 day on a medium without yucasin and transferred to a medium with different yucasin concentrations to allow for IZE survival. Treating IZE explants with yucasin from the beginning of the culture (without preculture) resulted in the death of the explants (Karami et al., 2023).

### Induction of *BDL*, *bdl*, and **Δ***ARF5* expression in transgenic plants

The action of BDL and bdl was controlled by adding dexamethasone (water-soluble DEX, Sigma; #D2915) to the media at a concentration of 10 or 30 µM. The expression of Δ*ARF5* was controlled by adding 1 µM β-estradiol (ethanol-soluble E, Sigma; # E8875) to the media.

### RNA isolation, reverse transcription, qualitative and quantitative PCR

RNA was isolated from different samples, depending on culture age, tissue from 300 (0, 1st day), 150 (5th day) to 50 (10th day) explants were used for RNA extraction. A NucleoSpin RNA Plant Kit (Macherey-Nagel) was used to isolate the RNA from the samples. The RNAs were treated with RQ1 RNase-free DNase I (Promega) to control DNA contamination according to the manufacturer’s instructions. The first-strand cDNA was produced from 1,000 ng of RNA using a RevertAid First Strand cDNA Synthesis Kit (Fermentas). The product of the reverse transcription was used for RT-PCR and RT-qPCR reactions to detect and quantify gene transcripts. Undiluted cDNA (1 µL) was used for RT-PCR. cDNA diluted with water at a 1:4 ratio (2.5 µL) was used for the RT-qPCR reactions. RT-qPCR was carried out in a 10 µL reaction volume using a LightCycler^®^ 480 SYBR^TM^ Green I Master (Roche) kit, a LightCycler^®^ 96 Multiwell Plate 96, and MultiwellSealing Foil (Roche). The relative RNA levels were calculated and normalized to the internal control of the AT4G27090 (*TIN*) gene encoding the 60S ribosomal protein (Thellin et al., 1999). Fold change values were calculated using the comparative 2^-ΔΔCt^ method, where ΔΔCt represents Δ*Ct* ^reference^ ^condition^ – Δ*Ct* ^compared^ ^condition^. The 40-Δ*Ct* method was used to compare the expression levels of *MP* and *MP11ir* (Czechowski et al., 2004). The plant tissues for gene expression analysis were produced in three biological replicates, and two technical replicates were analyzed. Primer sequences for *MP11ir*, *MP*, *TIN*, *TAA1*, *TARs*, and *YUCCAs* are in Suppl. Tab. S1.

### GUS staining

The GUS enzyme activity was performed as described by Bassuner et al. (2007). The explants were stained in a standard X-Gluc solution at 37 °C for 12 h. The experiments were repeated two times for each studied combination, and at least 30 somatic embryos were subjected to GUS signal detection in each repetition. The GUS staining pattern was analyzed using a microscope Zeiss Axioscope.A1 ZEN.

### ClearSee alpha clearing and Renaissance SR2200 staining

The explants were fixed using a fixative solution (4% paraformaldehyde in 1X PBS with 0.05% Triton-X100 (PBS-T)) and incubated in a 1.5 mL fixative in a microcentrifuge tube for one hour at room temperature. The tubes were covered with aluminum foil throughout the procedure to be light-tight. The samples were vacuum infiltrated for ∼ 30 min on ice to remove air bubbles. Then, the samples were incubated overnight at 4 °C with slow rotation. The following day, samples were washed three times with 1X PBS-T at room temperature for one hour each. A ClearSee alpha solution was prepared according to Kurihara et al. (2021). The ClearSee solution was supplemented with 50 mM sodium sulfite as an antioxidant. The clearing was extended to 7 days with gentle rotation (10-20 rpm) at room temperature to ensure thorough tissue penetration and clearing. After clearing, a staining step was performed by mixing a 0.1% (v/v) Renaissance SR2200 (Renaissance Chemicals, UK) solution in a ClearSee alpha solution (Attuluri et al., 2022). After staining, the samples were washed in a fresh ClearSee alpha solution for one hour and mounted on slides.

### Microscopy

Expression pattern analysis was performed on a ZEISS AxioZoom.V16 microscope equipped with an Apotome2 module, allowing for an overview of GFP signal localization. For higher resolution and subcellular localization, particularly to assess nuclear localization of the GFP signal, samples were subsequently examined using a ZEISS LSM 800 laser scanning confocal microscope with a correction collar 10 × magnification objective at 1024 × 1024 pixel resolution. Two laser lines were used: 405 nm for imaging Renaissance combined with 488 nm for GFP imaging. The images were captured in 12 bits. The laser intensity and detector gain were kept constant for all samples.

### Evaluation of indolic compound content

A colorimetric technique that enabled the detection of indolic compounds, including IAA, was applied (Bric et al., 1991; Wójcikowska et al., 2013). Each analysis was carried out in five biological replicates.

### Evaluation of IAA content

The concentration of indole-3-acetic acid (IAA) was evaluated in a 10-day-old IZE-culture of *pXVE::*Δ*ARF5 pDR5::GFP* transgenic line with 1 µM β-estradiol-induced Δ*ARF5* overexpression as described by Li-Marchetti et al. (2015). Fresh tissue was immediately frozen in liquid nitrogen and stored at −80 °C. The samples were freeze-dried and ground. Approximately 3 mg of dry powder was extracted for each sample with 0.8 mL of acetone/water/acetic acid (80/19/1 v:v:v). Indole-3-acetic acid stable labeled isotope used as internal standards were prepared, as described by Le Roux et al. (2014). Analysis was performed on four biological replicates.

### ARF binding regions

We employed the DAP-Seq profiles for the MP and ARF2 transcription factors (O’Malley et al., 2016) available at the PlantCistrome database (http://neomorph.salk.edu/PlantCistromeDB). We analyzed (−1500; +100) *TAA1/TAR* and *YUC* gene promoters and detected TGTCnn sequences and ARF-binding DAP-Seq peaks. We utilized a suite of R packages, including biomaRt, dplyr, and ggplot2, for computational analysis and data visualization.

### Statistical analysis

Phenotype scoring and quantification data were collected in EXCEL (Microsoft Corp.) and imported into STATISTICA (TIBCO Software Inc., Palo Alto, CA, USA) for statistical analysis and to generate graphs. For one-on-one comparisons, The Student’s T-test was used (*p* < 0.05). For intercomparison of more than two data points, the two-way ANOVA analysis (*p* < 0.05) followed by Tukey’s HSD test (*p* < 0.05, and *p* < 0.01) was used to determine any significant differences between the compared combinations. The graphs show the means with the standard deviations (SD).

## Data availability

Raw images are provided in the Zenodo repository (DOI: 10.5281/zenodo.15357086)

## Supplementary Data

Supplementary Figure S1. MP11ir is expressed during somatic embryogenesis.

Supplementary Figure S2. The induction of somatic embryogenesis depends on 2,4-D and NAA, not IAA.

Supplementary Figure S3. Expression level of auxin biosynthetic genes in *pXVE::*Δ*ARF5 pDR5::GFP* somatic embryos.

Supplementary Figure S4. Analysis of the *TAA1*, *TAR1*, and *YUC* promoters of MP binding sites.

Supplementary Figure S5. Expression pattern of TAA1, TAR1, TAR2, and YUC1-11 during somatic embryogenesis induction.

Supplementary Figure S6. Effects of repressing auxin biosynthesis in *pXVE::*Δ*ARF5 pDR5::GFP* during somatic embryogenesis induction.

Supplementary Table S1. The primers used in experiments.

## Funding

B.W. was supported by the Polish National Agency for Academic Exchange (BPN/BEK/2021/1/00278/U/00001). This work was supported by the project TowArds Next GENeration Crops, reg. no. CZ.02.1.01/00/0.0/22_008/0004581 of the ERDF Programme Johannes Amos Comenius from the Ministry of Education, Youth and Sports of the Czech Republic. The IJPB benefits from the support of Saclay Plant Sciences-SPS (ANR-17-EUR-0007). This research received funding from the National Science Centre, Poland, under the OPUS call in the Weave programme (2023/51/I/NZ1/00704) and the Czech Science Foundation (25-14425L).

## Acknowledgments

We acknowledge the CEITEC core facilities Plant Sciences and CELLIM, supported by the MEYS CR (LM2023050 Czech-BioImaging). This work has benefited from the support of IJPB’s Plant Observatory platforms P0-Chem. We are grateful to Lucia Colombo (Dipartimento di BioScienze, Università degli Studi di Milano, Milano, Italy), Dolf Weijers (Laboratory of Biochemistry, Wageningen University, and Research, Wageningen, The Netherlands), Bastiaan Bargmann (School of Plant and Environmental Sciences, Virginia Polytechnic Institute and State University, Blacksburg, USA) for providing seeds of Arabidopsis transgenic lines; Javier Brumos Institute for Plant Molecular and Cell Biology (IBMCP), CSIC-Universitat Politècnica de València, Spain, for consultation; Weronika Buchcik, Jagoda Polak, Wiktoria Remsak, Agnieszka Kiwior-Wesołowska for technical assistance.

## Author contributions

B.W. and H.R.B. designed and supervised the project. B.W., S.B., V.M., and S.C. conducted the experiments and analyzed the data. B.W. and H.R.B. wrote the manuscript. All authors discussed the results and approved the manuscript.

## Conflict of interest

The authors declare that they have no conflict of interest.

## Notes

https://doi.org/10.5281/zenodo.15357086

### Competing Interest Statement

The authors have declared no competing interest.

### Summary of Updates

One author was added for auxin levels measurement in Figure 8. Added data on gene expression in complemented plants Updated the expression patterns on MP and MP11ir.

## References

Alonso, J.M., Stepanova, A.N., Leisse, T.J., Kim, C.J., Chen, H., Shinn, P., Stevenson, D.K., Zimmerman, J., Barajas, P., Cheuk, R., Gadrinab, C., Heller, C., Jeske, A., Koesema, E., Meyers, C.C., Parker, H., Prednis, L., Ansari, Y., Choy, N., Deen, H., Geralt, M., Hazari, N., Hom, E., Karnes, M., Mulholland, C., Ndubaku, R., Schmidt, I., Guzman, P., Aguilar-Henonin, L., Schmid, M., Weigel, D., Carter, D.E., Marchand, T., Risseeuw, E., Brogden, D., Zeko, A., Crosby, W.L., Berry, C.C., Ecker, J.R., 2003. Genome-wide insertional mutagenesis of Arabidopsis thaliana. Science 301, 653–657.

Attuluri, V.P.S., Sánchez López, J.F., Maier, L., Paruch, K., Robert, H.S., 2022. Comparing the efficiency of six clearing methods in developing seeds of Arabidopsis thaliana. Plant Reprod 35, 279–293.

Awada, R., Lepelley, M., Breton, D., Charpagne, A., Campa, C., Berry, V., Georget, F., Breitler, J.C., Léran, S., Djerrab, D., Martinez-Seidel, F., Descombes, P., Crouzillat, D., Bertrand, B., Etienne, H., 2023. Global transcriptome profiling reveals differential regulatory, metabolic and hormonal networks during somatic embryogenesis in Coffea arabica. BMC Genomics 24, 41.

Bai, B., Su, Y.H., Yuan, J., Zhang, X.S., 2013. Induction of somatic embryos in Arabidopsis requires local YUCCA expression mediated by the down-regulation of ethylene biosynthesis. Mol Plant 6, 1247–1260.

Ballesteros, D., Martínez, M.T., Sánchez-Romero, C., Montalbán, I.A., Sales, E., Moncaleán, P., Arrillaga, I., Corredoira, E., 2024. Current status of the cryopreservation of embryogenic material of woody species. Frontiers in Plant Science 14, 1337152.

Banasiak, A., Biedroń, M., Dolzblasz, A., Berezowski, M.A., 2019. Ontogenetic Changes in Auxin Biosynthesis and Distribution Determine the Organogenic Activity of the Shoot Apical Meristem in pin1 Mutants. International Journal of Molecular Sciences 20, 180.

Bassuner, B.M., Lam, R., Lukowitz, W., Yeung, E.C., 2007. Auxin and root initiation in somatic embryos of Arabidopsis. Plant Cell Rep 26, 1–11.

Besnard, F., Refahi, Y., Morin, V., Marteaux, B., Brunoud, G., Chambrier, P., Rozier, F., Mirabet, V., Legrand, J., Lainé, S., 2014. Cytokinin signalling inhibitory fields provide robustness to phyllotaxis. Nature 505, 417–421.

Bhatia, N., Bozorg, B., Larsson, A., Ohno, C., Jönsson, H., Heisler, M.G., 2016. Auxin Acts through MONOPTEROS to Regulate Plant Cell Polarity and Pattern Phyllotaxis. Curr Biol 26, 3202–3208.

Boer, D.R., Freire-Rios, A., van den Berg, W.A., Saaki, T., Manfield, I.W., Kepinski, S., López-Vidrieo, I., Franco-Zorrilla, J.M., de Vries, S.C., Solano, R., Weijers, D., Coll, M., 2014. Structural basis for DNA binding specificity by the auxin-dependent ARF transcription factors. Cell 156, 577–589.

Boutilier, K., Offringa, R., Sharma, V.K., Kieft, H., Ouellet, T., Zhang, L., Hattori, J., Liu, C.M., van Lammeren, A.A., Miki, B.L., Custers, J.B., van Lookeren Campagne, M.M., 2002. Ectopic expression of BABY BOOM triggers a conversion from vegetative to embryonic growth. Plant Cell 14, 1737–1749.

Bric, J.M., Bostock, R.M., Silverstone, S.E., 1991. Rapid in situ assay for indoleacetic acid production by bacteria immobilized on a nitrocellulose membrane. Applied and environmental Microbiology 57, 535–538.

Brumos, J., Zhao, C., Gong, Y., Soriano, D., Patel, A.P., Perez-Amador, M.A., Stepanova, A.N., Alonso, J.M., 2019. An Improved Recombineering Toolset for Plants. The Plant Cell 32, 100–122.

Cancé, C., Martin-Arevalillo, R., Boubekeur, K., Dumas, R., 2022. Auxin response factors are keys to the many auxin doors. New Phytol 235, 402–419.

Cavalleri, A., Astori, C., Truskina, J., Cucinotta, M., Farcot, E., Chrysanthou, E., Xu, X., Muino, J.M., Kaufmann, K., Kater, M.M., Vernoux, T., Weijers, D., Bennett, M.J., Bhosale, R., Bishopp, A., Colombo, L., 2024. Auxin-dependent post-translational regulation of MONOPTEROS in the Arabidopsis root. Cell Reports 43(12).

Capote, T., Usié, A., Barbosa, P., Ramos, M., Morais-Cecílio, L., Gonçalves, S., 2019. Transcriptome dynamics of cork oak (Quercus suber) somatic embryogenesis reveals active gene players in transcription regulation and phytohormone homeostasis of embryo development. Tree Genetics & Genomes 15, 52.

Caumon, H., Vernoux, T., 2023. A matter of time: auxin signaling dynamics and the regulation of auxin responses during plant development. Journal of Experimental Botany 74, 3887–3902.

Chandler, J.W., 2016. Auxin response factors. Plant Cell Environ 39, 1014–1028.

Chen, Y., Xu, X., Liu, Z., Zhang, Z., XuHan, X., Lin, Y., Lai, Z., 2020. Global scale transcriptome analysis reveals differentially expressed genes involve in early somatic embryogenesis in Dimocarpus longan Lour. BMC Genomics 21, 4.

Chen, Z., Debernardi, J.M., Dubcovsky, J., Gallavotti, A., 2022. The combination of morphogenic regulators BABY BOOM and GRF-GIF improves maize transformation efficiency. bioRxiv, 2022.2009.2002.506370.

Ckurshumova, W., Berleth, T., 2015. Overcoming recalcitrance-Auxin response factor functions in plant regeneration. Plant signaling & behavior 10, e993293.

Cohen, J.D., Strader, L.C., 2024. An auxin research odyssey: 1989–2023. The Plant Cell 36, 1410–1428.

Cole, M., Chandler, J., Weijers, D., Jacobs, B., Comelli, P., Werr, W., 2009. DORNRÖSCHEN is a direct target of the auxin response factor MONOPTEROS in the Arabidopsis embryo. Development 136, 1643–1651.

Cucinotta, M., Cavalleri, A., Guazzotti, A., Astori, C., Manrique, S., Bombarely, A., Oliveto, S., Biffo, S., Weijers, D., Kater, M.M., Colombo, L., 2021. Alternative Splicing Generates a MONOPTEROS Isoform Required for Ovule Development. Curr Biol 31, 892–899.e893.

Czechowski, T., Bari, R.P., Stitt, M., Scheible, W.-R., Udvardi, M.K., 2004. Real-time RT-PCR profiling of over 1400 Arabidopsis transcription factors: unprecedented sensitivity reveals novel root- and shoot-specific genes. The Plant Journal 38, 366–379.

Elhiti, M., Stasolla, C., 2022. Transduction of Signals during Somatic Embryogenesis. Plants (Basel) 11.

Fambrini, M., Usai, G., Pugliesi, C., 2022. Induction of Somatic Embryogenesis in Plants: Different Players and Focus on WUSCHEL and WUS-RELATED HOMEOBOX (WOX) Transcription Factors. Int J Mol Sci 23.

Freire-Rios, A., Tanaka, K., Crespo, I., van der Wijk, E., Sizentsova, Y., Levitsky, V., Lindhoud, S., Fontana, M., Hohlbein, J., Boer, D.R., Mironova, V., Weijers, D., 2020. Architecture of DNA elements mediating ARF transcription factor binding and auxin-responsive gene expression in Arabidopsis. Proc Natl Acad Sci U S A 117, 24557–24566.

Gaj, M.D., 2001. Direct somatic embryogenesis as a rapid and efficient system for in vitro regeneration of Arabidopsis thaliana. Plant Cell, Tissue and Organ Culture 64, 39–46.

Gaj, M.D., Zhang, S., Harada, J.J., Lemaux, P.G., 2005. Leafy cotyledon genes are essential for induction of somatic embryogenesis of Arabidopsis. Planta 222, 977–988.

Gamborg, O.L.c., Miller, R.A., Ojima, K., 1968. Nutrient requirements of suspension cultures of soybean root cells. Experimental cell research 50, 151–158.

Garrett, J.J., Meents, M.J., Blackshaw, M.T., Blackshaw, L.C., Hou, H., Styranko, D.M., Kohalmi, S.E., Schultz, E.A., 2012. A novel, semi-dominant allele of MONOPTEROS provides insight into leaf initiation and vein pattern formation. Planta 236, 297–312.

Ge, X., Yu, X., Liu, Z., Yuan, J., Qin, A., Wang, Y., Chen, Y., Qin, W., Liu, Y., Liu, X., Zhou, Y., Wang, P., Yang, J., Liu, H., Zhao, Z., Hu, M., Zhang, Y., Sun, S., Herrera-Estrella, L., Phan Tran, L.S., Sun, X., Li, F., 2025. Spatiotemporal transcriptome and metabolome landscapes of cotton somatic embryos. Nature Communications, 16, 859.

Ghelli, R., Brunetti, P., Marzi, D., Cecchetti, V., Costantini, M., Lanzoni-Rossi, M., Scaglia Linhares, F., Costantino, P., Cardarelli, M., 2023. The full-length Auxin Response Factor 8 isoform ARF8.1 controls pollen cell wall formation and directly regulates TDF1, AMS and MS188 expression. Plant J 113, 851–865.

Ghelli, R., Brunetti, P., Napoli, N., De Paolis, A., Cecchetti, V., Tsuge, T., Serino, G., Matsui, M., Mele, G., Rinaldi, G., Palumbo, G.A., Barozzi, F., Costantino, P., Cardarelli, M., 2018. A Newly Identified Flower-Specific Splice Variant of AUXIN RESPONSE FACTOR8 Regulates Stamen Elongation and Endothecium Lignification in Arabidopsis. Plant Cell 30, 620–637.

Gliwicka, M., Nowak, K., Balazadeh, S., Mueller-Roeber, B., Gaj, M.D., 2013. Extensive modulation of the transcription factor transcriptome during somatic embryogenesis in Arabidopsis thaliana. PLoS One 8, e69261.

Gonzalez, J.H., Taylor, J.S., Reed, K.M., Wright, R.C., Bargmann, B.O., 2021. Temporal control of morphogenic factor expression determines efficacy in enhancing regeneration. Plants 10, 2271.

Guilfoyle, T.J., 2015. The PB1 domain in auxin response factor and Aux/IAA proteins: a versatile protein interaction module in the auxin response. Plant Cell 27, 33–43.

Guo, F., Wang, H., Lian, G., Cai, G., Liu, W., Zhang, H., Li, D., Zhou, C., Han, N., Zhu, M., Su, Y., Seo, P.J., Xu, L., Bian, H., 2023a. Initiation of scutellum-derived callus is regulated by an embryo-like developmental pathway in rice. Communications Biology 6, 457.

Guo, X., Wang, T., Jiang, L., Qi, H., Zhang, Z., 2023b. PlaASDB: a comprehensive database of plant alternative splicing events in response to stress. BMC plant biology 23, 225.

Harding, E.W., Tang, W., Nichols, K.W., Fernandez, D.E., Perry, S.E., 2003. Expression and maintenance of embryogenic potential is enhanced through constitutive expression of AGAMOUS-Like 15. Plant Physiol 133, 653–663.

Hardtke, C.S., Ckurshumova, W., Vidaurre, D.P., Singh, S.A., Stamatiou, G., Tiwari, S.B., Hagen, G., Guilfoyle, T.J., Berleth, T., 2004. Overlapping and non-redundant functions of the Arabidopsis auxin response factors MONOPTEROS and NONPHOTOTROPIC HYPOCOTYL 4. Development 131, 1089–1100.

Hasnain, A., Naqvi, S.A.H., Ayesha, S.I., Khalid, F., Ellahi, M., Iqbal, S., Hassan, M.Z., Abbas, A., Adamski, R., Markowska, D., 2022. Plants in vitro propagation with its applications in food, pharmaceuticals and cosmetic industries; current scenario and future approaches. Frontiers in plant science 13, 1009395.

Hu, Y., Zhou, L., Huang, M., He, X., Yang, Y., Liu, X., Li, Y., Hou, X., 2018. Gibberellins play an essential role in late embryogenesis of Arabidopsis. Nature Plants 4, 289–298.

Indoliya, Y., Tiwari, P., Chauhan, A.S., Goel, R., Shri, M., Bag, S.K., Chakrabarty, D., 2016. Decoding regulatory landscape of somatic embryogenesis reveals differential regulatory networks between japonica and indica rice subspecies. Sci Rep 6, 23050.

Juárez-González, V.T., López-Ruiz, B.A., Baldrich, P., Luján-Soto, E., Meyers, B.C., Dinkova, T.D., 2019. The explant developmental stage profoundly impacts small RNA-mediated regulation at the dedifferentiation step of maize somatic embryogenesis. Sci Rep 9, 14511.

Junker, A., Mönke, G., Rutten, T., Keilwagen, J., Seifert, M., Thi, T.M., Renou, J.P., Balzergue, S., Viehöver, P., Hähnel, U., Ludwig-Müller, J., Altschmied, L., Conrad, U., Weisshaar, B., Bäumlein, H., 2012. Elongation-related functions of LEAFY COTYLEDON1 during the development of Arabidopsis thaliana. Plant J 71, 427–442.

Kadokura, S., Sugimoto, K., Tarr, P., Suzuki, T., Matsunaga, S., 2018. Characterization of somatic embryogenesis initiated from the Arabidopsis shoot apex. Dev Biol 442, 13–27.

Karami, O., de Jong, H., Somovilla, V.J., Villanueva Acosta, B., Sugiarta, A.B., Ham, M., Khadem, A., Wennekes, T., Offringa, R., 2023. Structure–activity relationship of 2,4-D correlates auxinic activity with the induction of somatic embryogenesis in Arabidopsis thaliana. Plant J 116, 1355–1369.

Karami, O., Khadem, A., Rahimi, A., Zagari, N., Aigner, S. and Offringa, R., 2024. Transient efflux inhibition improves plant regeneration by natural auxins. Plant J 118, 295–303.

Karami, O., Philipsen, C., Rahimi, A., Nurillah, A.R., Boutilier, K., Offringa, R., 2023. Endogenous auxin maintains embryonic cell identity and promotes somatic embryo development in Arabidopsis. Plant J 113, 7–22.

Karami, O., Rahimi, A., Mak, P., Horstman, A., Boutilier, K., Compier, M., van der Zaal, B., Offringa, R., 2021. An Arabidopsis AT-hook motif nuclear protein mediates somatic embryogenesis and coinciding genome duplication. Nat Commun 12, 2508.

Kashkan, I., Timofeyenko, K., Růžička, K., 2022. How alternative splicing changes the properties of plant proteins. Quant Plant Biol 3, e14.

Klepikova, A.V., Kasianov, A.S., Gerasimov, E.S., Logacheva, M.D., Penin, A.A., 2016. A high resolution map of the Arabidopsis thaliana developmental transcriptome based on RNA seq profiling. The Plant Journal 88, 1058–1070.

Korasick, D.A., Westfall, C.S., Lee, S.G., Nanao, M.H., Dumas, R., Hagen, G., Guilfoyle, T.J., Jez, J.M., Strader, L.C., 2014. Molecular basis for AUXIN RESPONSE FACTOR protein interaction and the control of auxin response repression. Proc Natl Acad Sci U S A 111, 5427–5432.

Krasteva, G., Georgiev, V., Pavlov, A., 2021. Recent applications of plant cell culture technology in cosmetics and foods. Engineering in Life Sciences 21, 68–76.

Krogan, N.T., Ckurshumova, W., Marcos, D., Caragea, A.E., Berleth, T., 2012. Deletion of MP/ARF5 domains III and IV reveals a requirement for Aux/IAA regulation in Arabidopsis leaf vascular patterning. New Phytol 194, 391–401.

Krogan, N.T., Marcos, D., Weiner, A.I., Berleth, T., 2016. The auxin response factor MONOPTEROS controls meristem function and organogenesis in both the shoot and root through the direct regulation of PIN genes. New Phytol 212, 42–50.

Kurihara, D., Mizuta, Y., Nagahara, S., Higashiyama, T., 2021. ClearSeeAlpha: advanced optical clearing for whole-plant imaging. Plant and Cell Physiology 62, 1302–1310.

Lau, S., De Smet, I., Kolb, M., Meinhardt, H., Jürgens, G., 2011. Auxin triggers a genetic switch. Nat Cell Biol 13, 611–615.

Le Roux, C., Del Prete, S., Boutet-Mercey, S., Perreau, F., Balague, C., Roby, D., Fagard, M., Gaudin, V. 2014. The hnRNP-Q protein LIF2 participates in the plant immune response. PLoS ONE 9(6):e99343.

Ledwoń, A., Gaj, M.D., 2009. LEAFY COTYLEDON2 gene expression and auxin treatment in relation to embryogenic capacity of Arabidopsis somatic cells. Plant Cell Rep 28, 1677–1688.

Ledwoń, A., Gaj, M.D., 2011. LEAFY COTYLEDON1, FUSCA3 expression and auxin treatment in relation to somatic embryogenesis induction in Arabidopsis. Plant Growth Regulation 65, 157–167.

Lee, K., Kim, J.H., Park, O.S., Jung, Y.J., Seo, P.J., 2022. Ectopic expression of WOX5 promotes cytokinin signaling and de novo shoot regeneration. Plant Cell Rep 41, 2415–2422.

Lee, K., Wang, K., 2023. Strategies for genotype-flexible plant transformation. Current Opinion in Biotechnology 79, 102848.

Li, M., Wrobel-Marek, J., Heidmann, I., Horstman, A., Chen, B., Reis, R., Angenent, G.C., Boutilier, K., 2022. Auxin biosynthesis maintains embryo identity and growth during BABY BOOM-induced somatic embryogenesis. Plant Physiol 188, 1095–1110.

Li-Marchetti, C., Le Bras, C., Relion, D., Citerne, S., Huché-Thélier, L., Sakr, S., Morel, P., Crespel, L. 2015. Genotypic differences in architectural and physiological responses to water restriction in rose bush. Frontiers in Plant Science, 6, 355.

Liu, L., Yahaya, B.S., Li, J., Wu, F., 2024. Enigmatic role of auxin response factors in plant growth and stress tolerance. Frontiers in Plant Science 15, 1398818.

Liu, S., Zhang, Y., Feng, Q., Qin, L., Pan, C., Lamin-Samu, A.T., Lu, G., 2018a. Tomato AUXIN RESPONSE FACTOR 5 regulates fruit set and development via the mediation of auxin and gibberellin signaling. Sci Rep 8, 2971.

Liu, Z., Miao, L., Huo, R., Song, X., Johnson, C., Kong, L., Sundaresan, V., Yu, X., 2018b. ARF2–ARF4 and ARF5 are essential for female and male gametophyte development in Arabidopsis. Plant and Cell Physiology 59, 179–189.

Lotan, T., Ohto, M.-a., Yee, K.M., West, M.A., Lo, R., Kwong, R.W., Yamagishi, K., Fischer, R.L., Goldberg, R.B., Harada, J.J., 1998. Arabidopsis LEAFY COTYLEDON1 is sufficient to induce embryo development in vegetative cells. Cell 93, 1195–1205.

Ma, Y., Miotk, A., Šutiković, Z., Ermakova, O., Wenzl, C., Medzihradszky, A., Gaillochet, C., Forner, J., Utan, G., Brackmann, K., Galván-Ampudia, C.S., Vernoux, T., Greb, T., Lohmann, J.U., 2019. WUSCHEL acts as an auxin response rheostat to maintain apical stem cells in Arabidopsis. Nat Commun 10, 5093.

Ma, Y., Zhao, B., Zhang, D., Ouyang, K., Li, J., 2024. Transcriptomic and functional analyses identify genes associated with embryogenic callus formation in Neolamarckia cadamba. Industrial Crops and Products 216, 118686.

Martín, G., Márquez, Y., Mantica, F., Duque, P., Irimia, M., 2021. Alternative splicing landscapes in Arabidopsis thaliana across tissues and stress conditions highlight major functional differences with animals. Genome biology 22, 1–26.

Mattsson, J., Ckurshumova, W., Berleth, T., 2003. Auxin signaling in Arabidopsis leaf vascular development. Plant Physiol 131, 1327–1339.

McFarland, F.L., Kaeppler, H.F., 2024. History and current status of embryogenic culture based tissue culture, transformation and gene editing of maize (Zea mays L.). The Plant Genome, e20451.

Mehbub, H., Akter, A., Akter, M.A., Mandal, M.S.H., Hoque, M.A., Tuleja, M., Mehraj, H., 2022. Tissue Culture in Ornamentals: Cultivation Factors, Propagation Techniques, and Its Application. Plants (Basel) 11.

Meinke, D.W., 2020. Genome wide identification of EMBRYO DEFECTIVE (EMB) genes required for growth and development in Arabidopsis. New Phytologist 226, 306–325.

Mira, M.M., Day, S., Ibrahim, S., Hill, R.D., Stasolla, C., 2023. The Arabidopsis Phytoglobin 2 mediates phytochrome B (phyB) light signaling responses during somatic embryogenesis. Planta 257, 88.

Möller, B.K., Ten Hove, C.A., Xiang, D., Williams, N., López, L.G., Yoshida, S., Smit, M., Datla, R., Weijers, D., 2017. Auxin response cell-autonomously controls ground tissue initiation in the early Arabidopsis embryo. Proc Natl Acad Sci U S A 114, E2533–e2539.

Murashige, T., Skoog, F., 1962. A Revised Medium for Rapid Growth and Bio Assays with Tobacco Tissue Cultures. Physiologia Plantarum 15, 473–497.

Nanao, M.H., Vinos-Poyo, T., Brunoud, G., Thévenon, E., Mazzoleni, M., Mast, D., Lainé, S., Wang, S., Hagen, G., Li, H., Guilfoyle, T.J., Parcy, F., Vernoux, T., Dumas, R., 2014. Structural basis for oligomerization of auxin transcriptional regulators. Nat Commun 5, 3617.

O’Malley, R.C., Huang, S.C., Song, L., Lewsey, M.G., Bartlett, A., Nery, J.R., Galli, M., Gallavotti, A., Ecker, J.R., 2016. Cistrome and Epicistrome Features Shape the Regulatory DNA Landscape. Cell 165, 1280–1292.

Odat, O., Gardiner, J., Sawchuk, M.G., Verna, C., Donner, T.J., Scarpella, E., 2014. Characterization of an allelic series in the MONOPTEROS gene of Arabidopsis. Genesis 52, 127–133.

Paul, P., Joshi, S., Tian, R., Diogo Junior, R., Chakrabarti, M., Perry, S.E., 2022. The MADS-domain factor AGAMOUS-Like18 promotes somatic embryogenesis. Plant Physiol 188, 1617–1631.

Piya, S., Shrestha, S.K., Binder, B., Stewart, C.N., Jr., Hewezi, T., 2014. Protein-protein interaction and gene co-expression maps of ARFs and Aux/IAAs in Arabidopsis. Front Plant Sci 5, 744.

Powers, S.K., Holehouse, A.S., Korasick, D.A., Schreiber, K.H., Clark, N.M., Jing, H., Emenecker, R., Han, S., Tycksen, E., Hwang, I., Sozzani, R., Jez, J.M., Pappu, R.V., Strader, L.C., 2019. Nucleo-cytoplasmic Partitioning of ARF Proteins Controls Auxin Responses in Arabidopsis thaliana. Mol Cell 76, 177–190.e175.

Przemeck, G.K., Mattsson, J., Hardtke, C.S., Sung, Z.R., Berleth, T., 1996. Studies on the role of the Arabidopsis gene MONOPTEROS in vascular development and plant cell axialization. Planta 200, 229–237.

Qi, S., Zhao, R., Yan, J., Fan, Y., Huang, C., Li, H., Chen, S., Zhang, T., Kong, L., Zhao, J., Zhang, J., 2021. Global Transcriptome and Coexpression Network Analyses Reveal New Insights Into Somatic Embryogenesis in Hybrid Sweetgum (Liquidambar styraciflua × Liquidambar formosana). Front Plant Sci 12, 751866.

Quintana-Escobar, A.O., Nic-Can, G.I., Galaz Avalos, R.M., Loyola-Vargas, V.M., Gongora-Castillo, E., 2019. Transcriptome analysis of the induction of somatic embryogenesis in Coffea canephora and the participation of ARF and Aux/IAA genes. PeerJ 7, e7752.

Rademacher, E.H., Möller, B., Lokerse, A.S., Llavata-Peris, C.I., van den Berg, W., Weijers, D., 2011. A cellular expression map of the Arabidopsis AUXIN RESPONSE FACTOR gene family. Plant J 68, 597–606.

Robert, H.S., Grunewald, W., Sauer, M., Cannoot, B., Soriano, M., Swarup, R., Weijers, D., Bennett, M., Boutilier, K., Friml, J., 2015. Plant embryogenesis requires AUX/LAX-mediated auxin influx. Development 142, 702–711.

Sakamoto, Y., Kawamura, A., Suzuki, T., Segami, S., Maeshima, M., Polyn, S., De Veylder, L., Sugimoto, K., 2022. Transcriptional activation of auxin biosynthesis drives developmental reprogramming of differentiated cells. Plant Cell 34, 4348–4365.

Schlereth, A., Möller, B., Liu, W., Kientz, M., Flipse, J., Rademacher, E.H., Schmid, M., Jürgens, G., Weijers, D., 2010. MONOPTEROS controls embryonic root initiation by regulating a mobile transcription factor. Nature 464, 913–916.

Schulz, B., Segobye, K., 2016. 2, 4-D transport and herbicide resistance in weeds. Journal of Experimental Botany 67, 3177–3179.

Shi, X., Zhang, C., Liu, Q., Zhang, Z., Zheng, B., Bao, M., 2016. De novo comparative transcriptome analysis provides new insights into sucrose induced somatic embryogenesis in camphor tree (Cinnamomum camphora L.). BMC Genomics 17, 26.

Stone, S.L., Braybrook, S.A., Paula, S.L., Kwong, L.W., Meuser, J., Pelletier, J., Hsieh, T.F., Fischer, R.L., Goldberg, R.B., Harada, J.J., 2008. Arabidopsis LEAFY COTYLEDON2 induces maturation traits and auxin activity: Implications for somatic embryogenesis. Proc Natl Acad Sci U S A 105, 3151–3156.

Stone, S.L., Kwong, L.W., Yee, K.M., Pelletier, J., Lepiniec, L., Fischer, R.L., Goldberg, R.B., Harada, J.J., 2001. LEAFY COTYLEDON2 encodes a B3 domain transcription factor that induces embryo development. Proc Natl Acad Sci U S A 98, 11806–11811.

Su, Y.H., Liu, Y.B., Bai, B., Zhang, X.S., 2015. Establishment of embryonic shoot–root axis is involved in auxin and cytokinin response during Arabidopsis somatic embryogenesis. Frontiers in Plant Science 5, 123838.

Su, Y.H., Zhao, X.Y., Liu, Y.B., Zhang, C.L., O’Neill, S.D., Zhang, X.S., 2009. Auxin-induced WUS expression is essential for embryonic stem cell renewal during somatic embryogenesis in Arabidopsis. Plant J 59, 448–460.

Szécsi, J., Joly, C., Bordji, K., Varaud, E., Cock, J.M., Dumas, C., Bendahmane, M., 2006. BIGPETALp, a bHLH transcription factor is involved in the control of Arabidopsis petal size. The EMBO journal 25, 3912–3920.

Szyrajew, K., Bielewicz, D., Dolata, J., Wójcik, A.M., Nowak, K., Szczygieł-Sommer, A., Szweykowska-Kulinska, Z., Jarmolowski, A., Gaj, M.D., 2017. MicroRNAs Are Intensively Regulated during Induction of Somatic Embryogenesis in Arabidopsis. Front Plant Sci 8, 18.

Tanaka, M., Kikuchi, A., Kamada, H., 2008. The Arabidopsis histone deacetylases HDA6 and HDA19 contribute to the repression of embryonic properties after germination. Plant Physiology 146, 149–161.

Thakare, D., Tang, W., Hill, K., Perry, S.E., 2008. The MADS-domain transcriptional regulator AGAMOUS-LIKE15 promotes somatic embryo development in Arabidopsis and soybean. Plant Physiology 146, 1663–1672.

Thellin, O., Zorzi, W., Lakaye, B., De Borman, B., Coumans, B., Hennen, G., Grisar, T., Igout, A., Heinen, E., 1999. Housekeeping genes as internal standards: use and limits. Journal of Biotechnology 75, 291–295.

Thibaud-Nissen, F., Shealy, R.T., Khanna, A., Vodkin, L.O., 2003. Clustering of microarray data reveals transcript patterns associated with somatic embryogenesis in soybean. Plant Physiol 132, 118–136.

Tsugafune, S., Mashiguchi, K., Fukui, K., Takebayashi, Y., Nishimura, T., Sakai, T., Shimada, Y., Kasahara, H., Koshiba, T., Hayashi, K.-i., 2017. Yucasin DF, a potent and persistent inhibitor of auxin biosynthesis in plants. Scientific Reports 7, 13992.

Tvorogova, V.E., Fedorova, Y.A., Potsenkovskaya, E.A., Kudriashov, A.A., Efremova, E.P., Kvitkovskaya, V.A., Wolabu, T.W., Zhang, F., Tadege, M., Lutova, L.A., 2019. The WUSCHEL-related homeobox transcription factor MtWOX9-1 stimulates somatic embryogenesis in Medicago truncatula. Plant Cell, Tissue and Organ Culture (PCTOC) 138, 517–527.

Uc-Chuc, M.A., Pérez-Hernández, C., Galaz-Ávalos, R.M., Brito-Argaez, L., Aguilar-Hernández, V., Loyola-Vargas, V.M., 2020. YUCCA-Mediated Biosynthesis of the Auxin IAA Is Required during the Somatic Embryogenic Induction Process in Coffea canephora. Int J Mol Sci 21.

Varaud, E., Brioudes, F., Szécsi, J., Leroux, J., Brown, S., Perrot-Rechenmann, C., Bendahmane, M., 2011. AUXIN RESPONSE FACTOR8 regulates Arabidopsis petal growth by interacting with the bHLH transcription factor BIGPETALp. Plant Cell 23, 973–983.

Wang, F.X., Shang, G.D., Wu, L.Y., Xu, Z.G., Zhao, X.Y., Wang, J.W., 2020. Chromatin Accessibility Dynamics and a Hierarchical Transcriptional Regulatory Network Structure for Plant Somatic Embryogenesis. Dev Cell 54, 742–757.e748.

Wickramasuriya, A.M., Dunwell, J.M., 2015. Global scale transcriptome analysis of Arabidopsis embryogenesis in vitro. BMC Genomics 16, 301.

Wójcik, A.M., Wójcikowska, B., Gaj, M.D., 2020. Current Perspectives on the Auxin-Mediated Genetic Network that Controls the Induction of Somatic Embryogenesis in Plants. Int J Mol Sci 21.

Wójcikowska, B., Belaidi, S., Robert, H.S., 2023. Game of thrones among AUXIN RESPONSE FACTORs—over 30 years of MONOPTEROS research. Journal of Experimental Botany 74, 6904–6921.

Wójcikowska, B., Botor, M., Morończyk, J., Wójcik, A.M., Nodzyński, T., Karcz, J., Gaj, M.D., 2018. Trichostatin A Triggers an Embryogenic Transition in Arabidopsis Explants via an Auxin-Related Pathway. Front Plant Sci 9, 1353.

Wójcikowska, B., Gaj, M.D., 2015. LEAFY COTYLEDON2-mediated control of the endogenous hormone content: implications for the induction of somatic embryogenesis in Arabidopsis. Plant Cell, Tissue and Organ Culture (PCTOC) 121, 255–258.

Wójcikowska, B., Gaj, M.D., 2017. Expression profiling of AUXIN RESPONSE FACTOR genes during somatic embryogenesis induction in Arabidopsis. Plant Cell Rep 36, 843–858.

Wójcikowska, B., Jaskóła, K., Gąsiorek, P., Meus, M., Nowak, K., Gaj, M.D., 2013. LEAFY COTYLEDON2 (LEC2) promotes embryogenic induction in somatic tissues of Arabidopsis, via YUCCA-mediated auxin biosynthesis. Planta 238, 425–440.

Wu, L.Y., Shang, G.D., Wang, F.X., Gao, J., Wan, M.C., Xu, Z.G., Wang, J.W., 2022. Dynamic chromatin state profiling reveals regulatory roles of auxin and cytokinin in shoot regeneration. Dev Cell 57, 526–542.e527.

Wu, M.F., Yamaguchi, N., Xiao, J., Bargmann, B., Estelle, M., Sang, Y., Wagner, D., 2015. Auxin-regulated chromatin switch directs acquisition of flower primordium founder fate. Elife 4, e09269.

Xie, M., Huang, L., Song, L., O’Neil, R., Lewsey, M., Chen, H., Chen, H., Zhuo, R., Shokhirev, M., Alonso, J., Ecker, J., 2022. Defining in vivo transcriptional responses to auxin. Research Square.

Xu, C., Shen, Y., He, F., Fu, X., Yu, H., Lu, W., Li, Y., Li, C., Fan, D., Wang, H.C., 2019. Auxin mediated Aux/IAA ARF HB signaling cascade regulates secondary xylem development in Populus. New Phytologist 222, 752–767.

Xu, L., Liu, Y., Zhang, J., Wu, W., Hao, Z., He, S., Li, Y., Shi, J., Chen, J., 2024. Genomic survey and expression analysis of LcARFs reveal multiple functions to somatic embryogenesis in Liriodendron. BMC Plant Biology 24, 94.

Yang, X., Lin, Q., Udayabhanu, J., Hua, Y., Dai, X., Xin, S., Wang, X., Huang, H., Huang, T., 2024. An optimized somatic embryo transformation system assisted homozygous edited rubber tree generation method mediated by CRISPR/Cas9. bioRxiv, 2024.2003. 2014.585007.

Yue, P., Lu, Q., Liu, Z., Lv, T., Li, X., Bu, H., Liu, W., Xu, Y., Yuan, H., Wang, A., 2020. Auxin activated MdARF5 induces the expression of ethylene biosynthetic genes to initiate apple fruit ripening. New phytologist 226, 1781–1795.

Zeng, J., Deng, Y., Iqbal, S., Zhang, J., Wu, K., Ma, G., Li, L., Dai, G., Deng, R., Fang, L., Zeng, S., 2024. Direct somatic embryogenesis and related gene expression networks in leaf explants of Hippeastrum ‘Bangkok Rose’. Horticultural Plant Journal 10, 556–572.

Zhang, M.M., Zhang, H.K., Zhai, J.F., Zhang, X.S., Sang, Y.L., Cheng, Z.J., 2021. ARF4 regulates shoot regeneration through coordination with ARF5 and IAA12. Plant Cell Rep 40, 315–325.

Zheng, Q., Perry, S.E., 2014. Alterations in the transcriptome of soybean in response to enhanced somatic embryogenesis promoted by orthologs of Agamous-like15 and Agamous-like18. Plant Physiol 164, 1365–1377.

Zheng, Y., Ren, N., Wang, H., Stromberg, A.J., Perry, S.E., 2009. Global identification of targets of the Arabidopsis MADS domain protein AGAMOUS-Like15. The Plant Cell 21, 2563–2577.

Zhu, X., Xu, Z., Wang, G., Cong, Y., Yu, L., Jia, R., Qin, Y., Zhang, G., Li, B., Yuan, D., Tu, L., Yang, X., Lindsey, K., Zhang, X., Jin, S., 2023. Single-cell resolution analysis reveals the preparation for reprogramming the fate of stem cell niche in cotton lateral meristem. Genome Biol 24, 194.

