## Supplemental Table and Figures for "*MONOPTEROS* isoform *MP11ir* role during somatic embryogenesis in *Arabidopsis thaliana*"

**Supplementary Table S1.** **The primers used in experiments.**

| **Primers used in RT-qPCR/PCR analysis of *TF* and *YUC* genes** | | | |  |
| --- | --- | --- | --- | --- |
| **Gene** | **ID** | **pF 5’-3’** | **pR 5’-3’** | **Publication** |
| *MP11ir* | *AT1G19850* | TGGGTAATGTTTTGACTTGG | CCACAAACTCTTCCCATGGAT | Cucinotta et al., 2021 |
| *MP* | *AT1G19850* | CACTAAGGTTCAAAAAACCG | TTACGCATCCCACAAACTCTTCC | Wójcikowska and Gaj, 2017 |
| *TIN* | At4g27090 | GTCGTTATCGTCGACGTTGTT | CCTCGATCAAAGCCTTCTTCT | Wójcikowska et al., 2013 |
| *TAA1* | *AT1G70560* | TTCGTGGTCAATCTGGATCATGG | ACCACGTATCGTCACCGTACAC |  |
| *TAR1* | *AT1G23320* | CGCAGCGGTTCCTTATTACTCCAC | TTGTCGAACGTCCTTGCGTCTC | Present work |
| *TAR2* | *AT4G24670* | GCTCTTCACTGCTTCAAAGAGCAC | TCTGTCTTTCACCAAAGCCCATCC |  |
| *YUC1* | *At4g32540* | CGGAACACCGTTCATGTGT | CCGGTGACATTTTTCAGCTC | Wójcikowska et al., 2013 |
| *YUC2* | *At4g13260* | TTGTGGTTCGTGACTCGGTA | TTCAAGAGGGCCAAGTTTTG |  |
| *YUC3* | *At1g04610* | GATGGCCGTGTTCTTGAGAT | TCATGAGCCACACTCATAGC |  |
| *YUC4* | *At5g11320* | AACTCCCGTTCTTGATGTCG | AAAAACTATTCTCCTTAAGCCAATC |  |
| *YUC5* | *At5g43890* | TGTCCAGTCTGCTCGATACG | TTCTCGCCGGATTTGTACTC |  |
| *YUC6* | *At5g25620* | GGTAAAACTCCGGTTCTCGAC | TTGGAAATCCATCTTTCTTACTAAAC |  |
| *YUC7* | *At2g33230* | TGAAGAACACCGCAGGTAAA | CCAAGTCGTTTTCCTTAAGCC |  |
| *YUC8* | *At4g28720* | CGTCTCAAGCTTCACCTTCC | AGCCACTGGTCTCATCGAAC |  |
| *YUC9* | *At1g04180* | TGGTCGTTAGAAGCTCGGTT | CGGCGTCTTTCCTGTCAT |  |
| *YUC10* | *At1g48910* | TTACCGGAAAAGCTCCTGTC | TCACGTATTCATAGTCCTCTAACCA |  |
| *YUC11* | *At1g21430* | GAGAATGGCGAAGGTGTGAT | TAACACGTGCACCTGGCTAC |  |

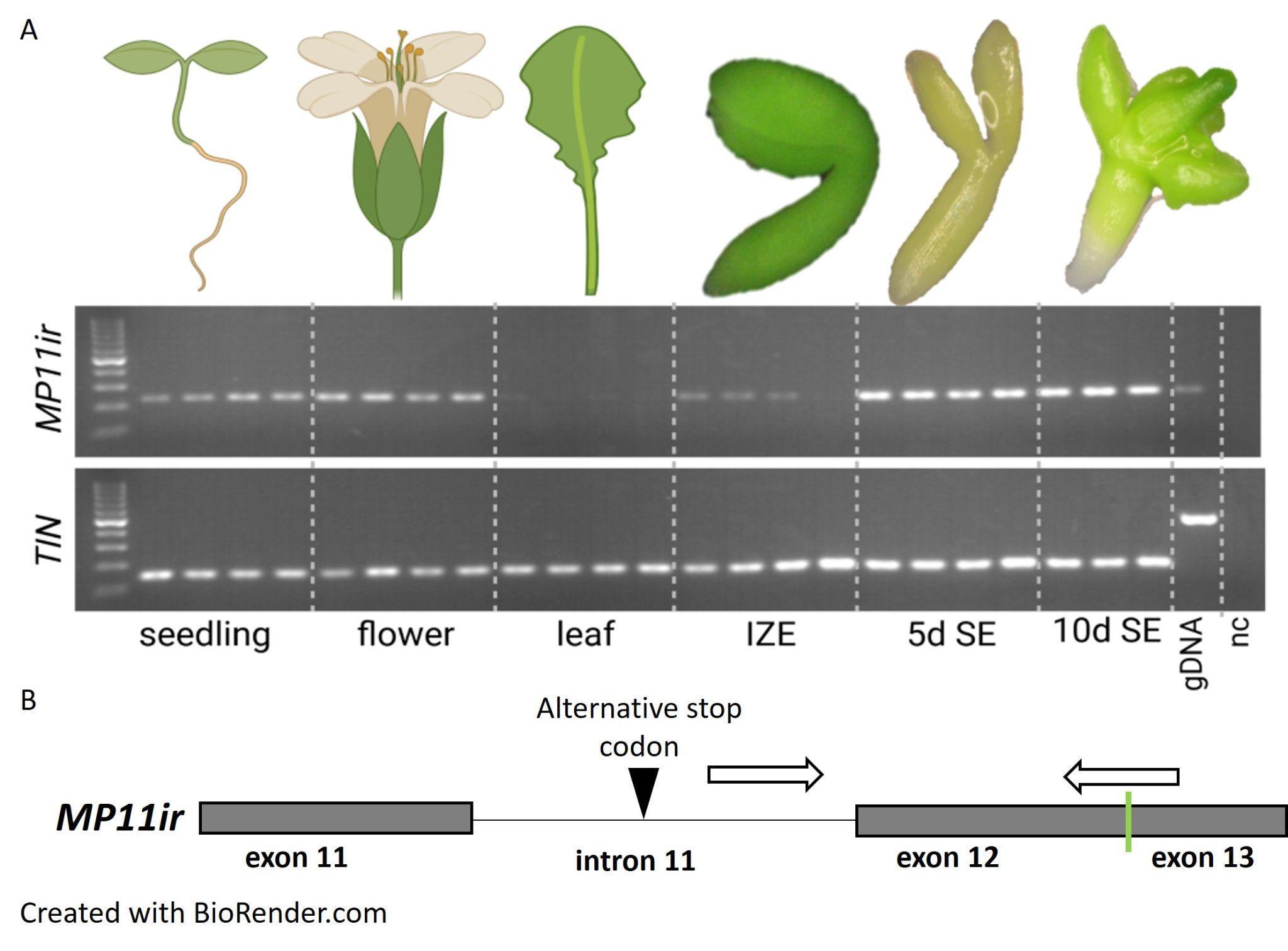

**Supplementary Figure S1. *MP11ir* is expressed during somatic embryogenesis.**

(**A**) Presence of *MP11ir* transcript during plant development *in vivo* and auxin-dependent SE induction *in vitro* (5.0 µM 2,4-D). (**B**) Location of primers used for detecting the *MP11ir* transcript. IZE – immature zygotic embryo; SE – somatic embryogenesis; 5d – 5th day of SE induction; 10d – 10th day of SE induction; gDNA – genomic DNA; nc – negative control; *TIN* – reference gene. Created with BioRender.com.

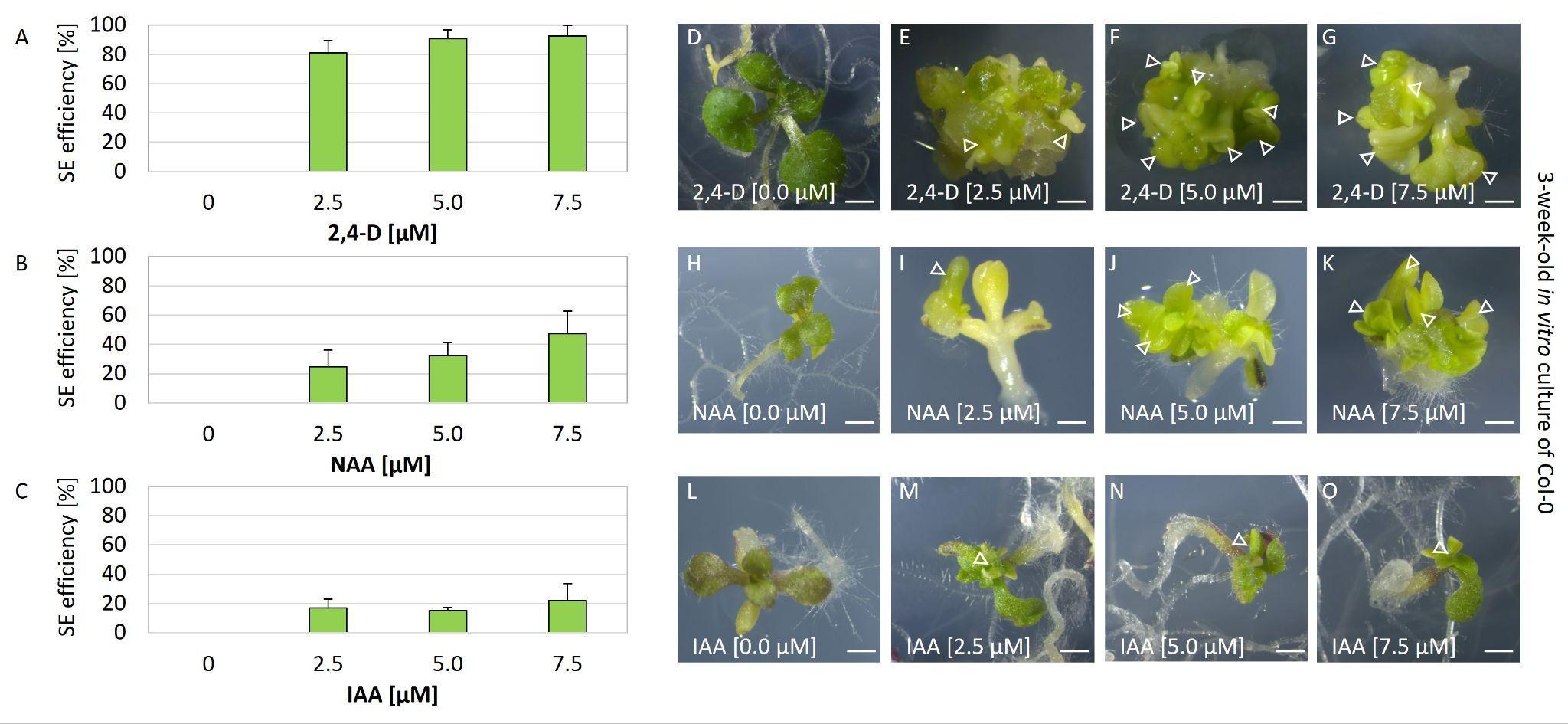

**Supplementary Figure S2. The induction of somatic embryogenesis depends on 2,4-D and NAA, not IAA.**

The capacity of IZE explants cultured for three weeks on media with different 2,4-D (**A, D-G**), NAA (**B, H-K**), and IAA (**C, L-O**) concentrations: 0.0 (**D, H, L**), 2.5 (**E, I, M**), 5.0 (**F, J, N**), and 7.5 µM (**G, K, O**); (n = 3; means ± SD are presented). IZE explants cultured on auxin-free medium developed seedlings after three weeks of culture (**D, H, L**). The increasing embryogenic capacity of the Col-0 IZE explants cultured for three weeks on media with different 2,4-D or NAA concentrations (2.5 (**E, I**), 5.0 (**F, J**), and 7.5 µM (**G, K**)). The weak embryogenic capacity of the Col-0 IZE explants cultured for three weeks on media with different IAA concentrations (2.5 (**M**), 5.0 (**N**), and 7.5 µM (**O**)). The white arrowhead indicates the somatic embryo. Size bars indicate 1 mm.

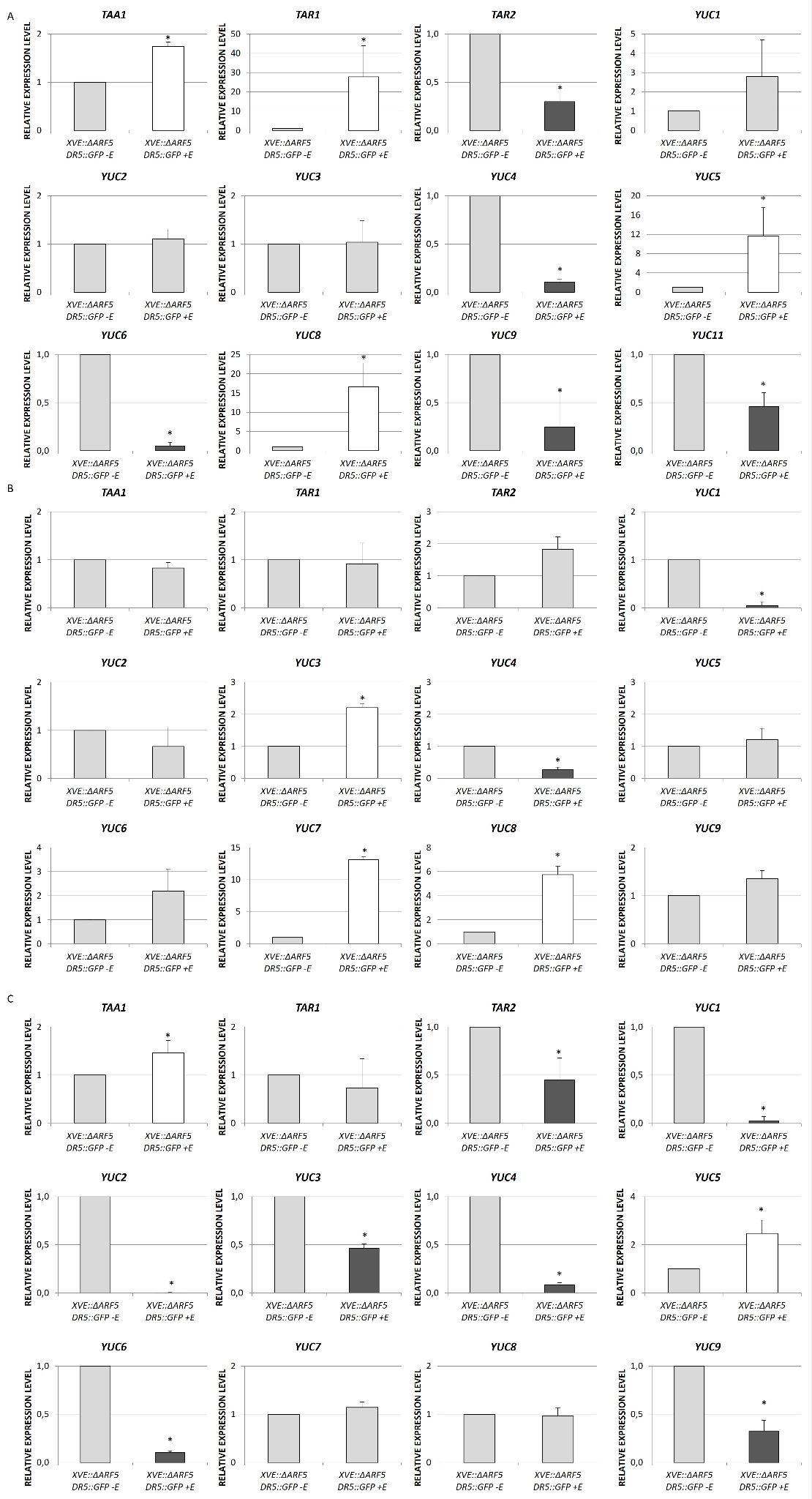

**Supplementary Figure S3. Expression level of auxin biosynthetic genes in *pXVE::ΔARF5 pDR5::GFP* somatic embryos.**

(**A-C**) Relative expression level of auxin biosynthesis genes in the 1- (**A**), 5- (**B**), and 10-day-old (**C**) embryogenic cultures of the *pXVE::ΔARF5 pDR5::GFP* transgenic line treated with 1 µM β-estradiol (+E). The relative expression level was normalized to internal control (At4g27090) and calibrated to the *pXVE::ΔARF5 pDR5::GFP* culture of the same age and untreated with 1 µM β-estradiol (-E). Value significantly different from the *pXVE::ΔARF5 pDR5::GFP* (-E) culture of the same age is marked with asterisks (**p* < 0.05; n = 3; means ± SD are presented). Gray bars indicate no statistical differences, white bars indicate that the gene is upregulated when *ΔARF5* is induced, and black bars when the gene expression is reduced.

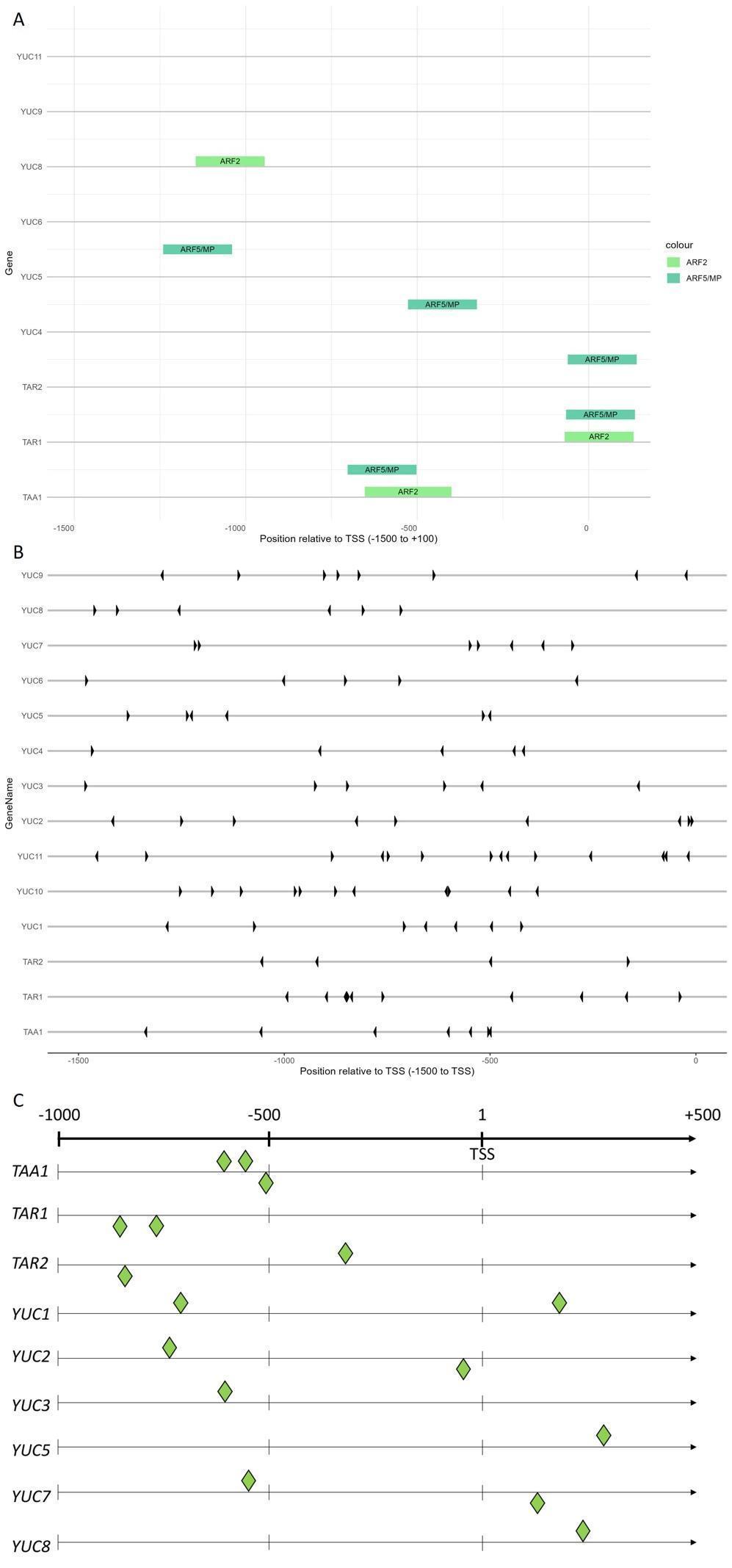

**Supplementary Figure S4. Analysis of the *TAA1*, *TAR1*, and *YUC* promoters for MP binding sites.**

(**A**) The *TAA1*, *TAR*, and *YUC* gene promoters were analyzed for MP and ARF2 binding sites. (**B**) Binding regions detected by DAP-seq in (O'Malley et al., 2016) TGTCnn sequences. (**C**) The presence of AuxRe element based on the PlantPan 4.0 database.

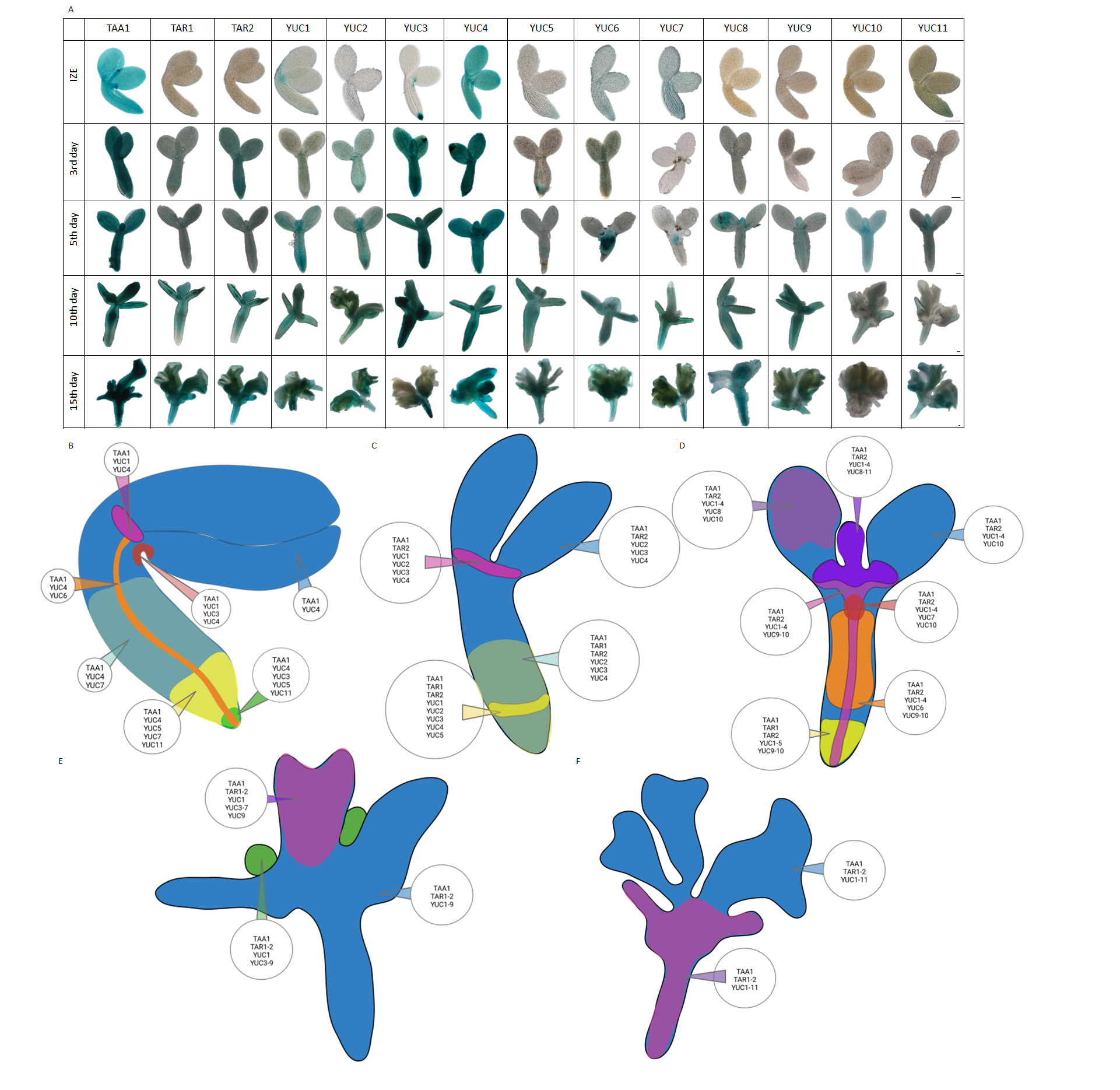

**Supplementary Figure S5.** **Expression pattern of TAA1, TAR1, TAR2, and YUC1-11 during somatic embryogenesis induction.**

(**A**) Spatio-temporal localization of TAA1, TAR1, TAR2, and YUC1-11 enzymes during SE process under *in vitro* culture. (**B-F**) Map of TAA1, TAR1, TAR2, YUC1-11 enzymes involved in auxin biosynthesis in IZE explants of reporter lines (**B**), and explants cultured on E5 medium and on 3- (**C**), 5- (**D**), 10- (**E**), 15-day-old (**F**) *in vitro* culture. Size bars indicate 100 μm (**A**).

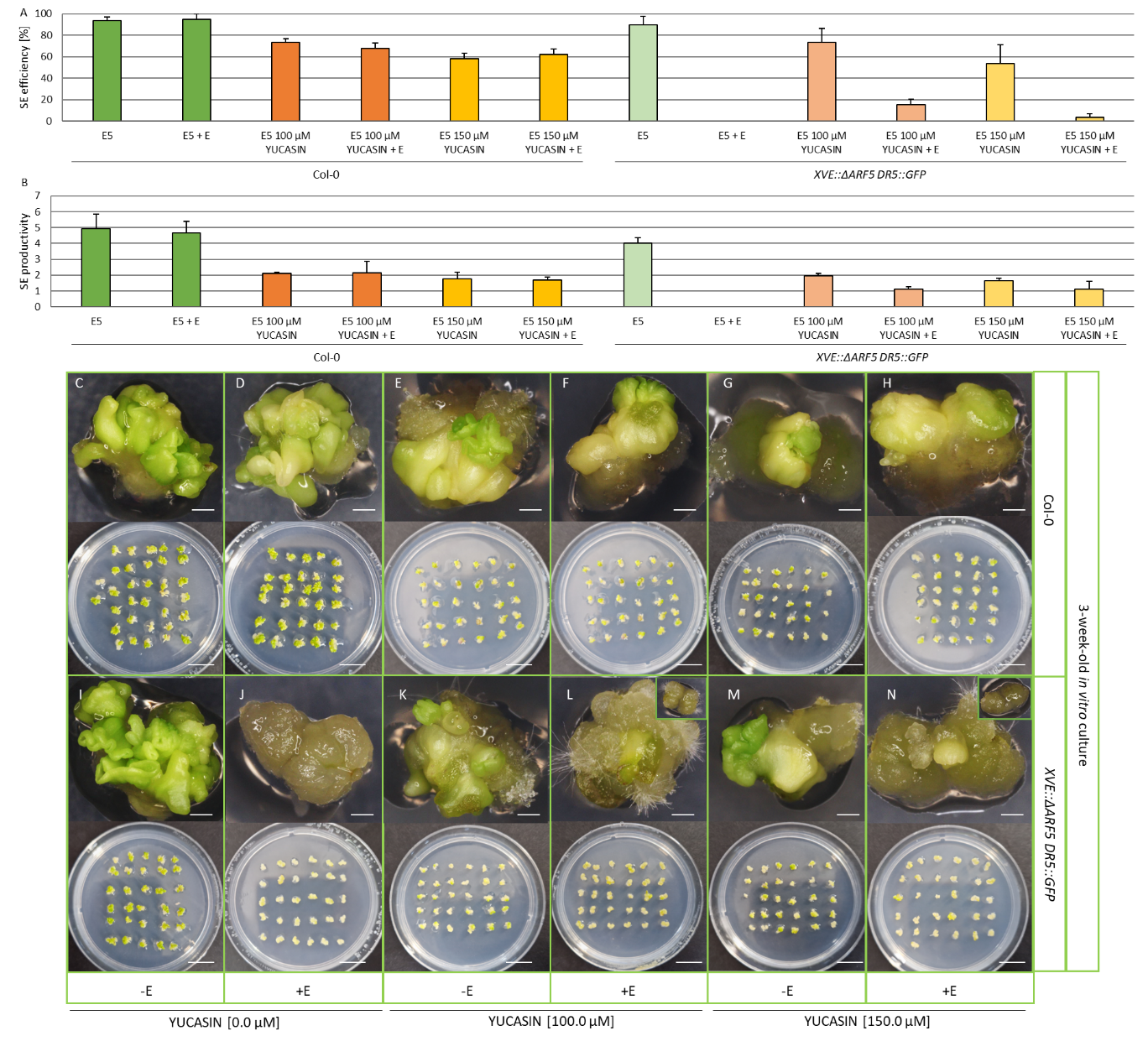
**Supplementary Figure S6. Effects of repressing auxin biosynthesis in *pXVE::ΔARF5 pDR5::GFP* during somatic embryogenesis induction.**

The effect of auxin biosynthesis inhibitor yucasin on the SE efficiency (**A**) and productivity (**B**) of the Col-0 (**C-H**) and *pXVE::ΔARF5 pDR5::GFP* (**I-N**) IZE explants. The capacity of Col-0 and *pXVE::ΔARF5 pDR5::GFP* IZE explants cultured for three weeks on E5 media with different yucasin concentrations: 0 (**C, D, I, J**), 100 (**E, F, K, L**), and 150 µM(**G, H, M, N**), supplemented (**D, F, H, J, L, N**) or not (**C, I, E, K, G, M**) with 1 µM estradiol (+E) for induction of *ΔARF5* overexpression; (n = 3; means ± SD are presented). Col-0 IZE explants cultured on control media (E5 and E5 + E) developed numerous somatic embryos after three weeks of culture (**C, D**). Different yucasin concentrations resulted in the decreasing embryogenic capacity of the Col-0 IZE explants cultured for three weeks on E5 and E5 + E media (**A, B, E-H**). The embryogenic capacity of the *pXVE::ΔARF5 pDR5::GFP* IZE explants cultured for three weeks on E5 media (**A, B, I**) is comparable to Col-0 IZE explants (**C**). Explants with *ΔARF5*-overexpression (E5 + E) lack a SE response (**J**)*.* Different yucasin concentrations resulted in the decreasing embryogenic capacity of the *pXVE::ΔARF5 pDR5::GFP* IZE explants cultured for three weeks on E5 media with (**A, B, K, M**). Explants overexpressing *ΔARF5* (+E) cultured on E5 media supplemented with yucasin at different concentrations were capable, but with very low efficiency and productivity, to regenerate somatic embryos. They mostly produced callus (**A, B, L, N**). Size bars indicate 1 mm for explant images and 1 cm for plate images.
